# Targeting a macrophage stemness factor to mitigate diseases post respiratory viral infection

**DOI:** 10.1101/2025.10.03.680366

**Authors:** Mohd Arish, Arka Sen Chaudhuri, Janli Tang, Harish Narasimhan, An Fang, Jinyi Tang, Xiaoqin Wei, Chaofan Li, In Su Cheon, Wei Qian, Farha Naz, Gislane Almeida-Santos, Samuel Patrick Young, Tanyalak Parimon, Yun Michael Shim, Robert Vassallo, Peter Chen, Jie Sun

## Abstract

Tissue-resident alveolar macrophages (AMs) rely on intrinsic stem-like programs for self-renewal and maintenance, yet the transcriptional networks that support these functions and their relevance to post-viral lung disease remain largely unknown. Here, we identify TCF4 (*Tcf7l2*) as a critical transcription factor that governs AM maturation and stemness. Loss of TCF4 impaired AM proliferation, shifted their identity toward a pro-inflammatory phenotype, and exacerbated host morbidity following influenza or SARS-CoV-2 infection. Conversely, enforced TCF4 expression promoted the expansion of mature AMs, and supported lung recovery, thereby protecting against severe acute viral disease. Mechanistically, TCF4 antagonized β-catenin-driven inflammatory transcription while preserving oxidative phosphorylation, defining a reciprocal regulatory axis essential for AM function. Notably, respiratory viral infections and exuberant interferon signaling suppressed TCF4 expression, which remains chronically reduced in murine and human lungs with post-COVID fibrosis. This downregulation is associated with persistent KRT8^hi^ dysplastic epithelium and collagen deposition. Moreover, aging diminished TCF4 levels and enforced TCF4 expression dampened age-associated decline of AM self-renewal. Furthermore, *in vivo* TCF4 overexpression after viral clearance enhanced mature AM accumulation, promoted lung epithelium regeneration, attenuated chronic tissue fibrosis and restored pulmonary physiologial function in aged lungs in a model of persistent pulmonary fibrosis post-acute viral infection. These findings have established TCF4 as a key regulator of AM stemness and identified a promising therapeutic target for long COVID and related chronic lung diseases through the modulation of embryonic-derived macrophage regenerative capacity by targeting TCF4.

## Introduction

Tissue-resident macrophages (RTMs) are recognized as long-lived, self-renewing sentinels that seed developing organs during embryogenesis (*1*). RTMs can persist throughout the adult life during homeostasis through controlled proliferation, reflecting the key hallmark of stemness. In particular, alveolar macrophages (AMs), a specialized subset of lung macrophages, have the great ability to proliferate and self-renew *in vitro* and *in vivo* (*2, 3*). This capacity for self-maintenance enables their long-term persistence in the lung with moderate contribution from monocytes during homeostasis throughout the adult life (*4*). AMs originate from fetal liver or yolk sac progenitors that populate the lungs during early development, and their population is sustained through signals such as GM-CSF, TGF-β, and neutrophil derived 12-HETE etc (*5–8*). However, the transcriptional mechanisms governing AM “stemness” remain incompletely understood. Furthermore, AMs play essential roles in host defense and recovery from respiratory infections (*9, 10*). Evidence has linked AM maintenance and self-renewal with better outcome after infection (*11*). To this end, macrophage proliferation is uniquely associated with a gene program that facilitates tissue repair (*12*), indicating a possibility of enhancing macrophage stemness to facilitate functional recovery after tissue injury. Nevertheless, it is still unknown currently whether we can target the AM self-renewal program to augment host resistance to infection and/or to facilitate tissue regeneration after acute infection.

Respiratory viral infections, such as influenza and SARS-CoV-2, are significant global health concerns due to their high transmissibility and potential for severe outcomes. Influenza, caused by influenza A and B viruses, leads to annual epidemics characterized by symptoms ranging from mild respiratory illness to severe complications like pneumonia. Seasonal influenza is associated with an average of more than 300 thousand respiratory deaths globally each year, among these about 67% were individuals of 65 years and older (*13*). COVID-19, caused by the SARS-CoV-2 virus, emerged in late 2019 and rapidly escalated into a global pandemic and killed for more than 20 millions of individuals including undocumented mortality (*14*). Moreover, the long-term persistence of chronic sequelae has been documented in cases of both influenza and COVID-19 (*15, 16*). Currently, over 60 million people worldwide are experiencing post-acute sequelae of SARS-CoV-2 infection (PASC, or long COVID), marked by persistent, recurrent, or new symptoms that emerge after the acute infection has been resolved (*17*). Recent studies have indicated the roles of aberrant immune responses in contribution to post-viral respiratory sequelae (*18–20*). Furthermore, an upregulation of gene programs associated with recruited monocyte-derived macrophages (MoAM), along with a downregulation of resident AM gene signatures, has been observed in lungs exhibiting persistent pathological responses following acute COVID-19 (*18, 21, 22*). These findings suggest that impaired repopulation of mature resident AMs may contribute to chronic lung conditions after acute infection. However, it remains unclear whether enhancing resident AM repopulation through the augmentation of AM proliferation and/or the promotion of maturation of MoAM towards resident AM can mitigate these outcomes.

The T-cell factor/lymphoid enhancer-binding factor (TCF/LEF) family of transcription factors plays a crucial role in stem cell function (*23, 24*). This family consists of four nuclear factors: TCF1, LEF1, TCF3 and TCF4. These factors serve as key downstream mediators of the Wnt/β-catenin signaling pathway, which is vital for regulating various cellular processes (*25*). In T cell lineages, TCF1 (encoded by *Tcf7*) is crucial for the self-renewal of “stem-like” CD8^+^ T cells generated in response to viral or tumor antigens. These stem-like cells are crucial for maintaining T cell immunity during prolonged antigen exposure (*26*). In CD4^+^ T cells, TCF1 plays a key role in regulating both stemness and follicular helper cell (Tfh) differentiation pathways, particularly in the context of autoimmunity. Currently, the roles of TCF family members in regulating macrophage self-renewal and stemness are unknown.

In this study, we identified TCF4 (encoded by *Tcf7l2*) as highly expressed in AMs and essential for their maturation and host defense against respiratory viral infections. Constitutive deletion of TCF4 disrupted AM development and maturation, while inducible ablation impaired their self-renewal and maintenance *in vivo*. Mechanistically, TCF4 antagonized β-catenin to support mitochondrial metabolism and restrain inflammation. Severe influenza and SARS-CoV-2 infections suppressed TCF4 expression *via* interferon signaling. Strikingly, TCF4 remained persistently downregulated in AMs from animal models of post-acute SARS-CoV-2 sequelae and in patients with persistent lung fibrosis following acute COVID-19. Myeloid-specific TCF4 deletion led to sustained transitional epithelial cells (KRT8^hi^) and chronic lung inflammation and fibrosis after acute infection. Conversely, enforced TCF4 expression enhanced AM self-renewal *in vitro* and *in vivo*, protected mice from lethal infection, and reduced acute lung injury. TCF4 expression also declined with age, correlating with age-associated reduction of AM stemness, while its overexpression promoted AM maturation and proliferation during aging, rescued age-related susceptibility to severe acute infection and mitigated chronic lung sequelae after the resolution of primary infection. Together, our findings reveal a central role for TCF4 in maintaining AM stemness and highlight its therapeutic potential for improving outcomes in acute and chronic lung disease, including long COVID.

## Results

### TCF4 modulates AM development and maturation

To determine the potential roles of TCF/LEF family of transcription factors AM stemness, we initially examined their expression in T cells, AMs, and peritoneal macrophages (PMs). Compared to T cells and PMs, AMs highly expressed TCF4 (Tcf7l2), with little to no expression of other TCF family members (Fig. 1A). Furthermore, both mouse and human AMs highly expressed TCF4 at the mRNA level (Fig. S1A-B). Additionally, other myeloid cells, including neutrophils and monocytes, exhibited minimal TCF4 expression compared to AMs (Fig. S1C). Given the key roles of GM-CSF and TGF-β in AM development and maturation (*7, 8, 27*), we tested whether GM-CSF and/or TGF-β stimulation alters TCF4 expression and found that combined stimulation with GM-CSF and TGF-β induced robust TCF4 expression in *in vitro* cultured AMs (Fig. 1B, S1D-E). Furthermore, GM-CSF plus TGF-β also induced TCF4 expression in bone marrow-derived monocytes (Fig 1C, S1F-H).

**Fig. 1.**
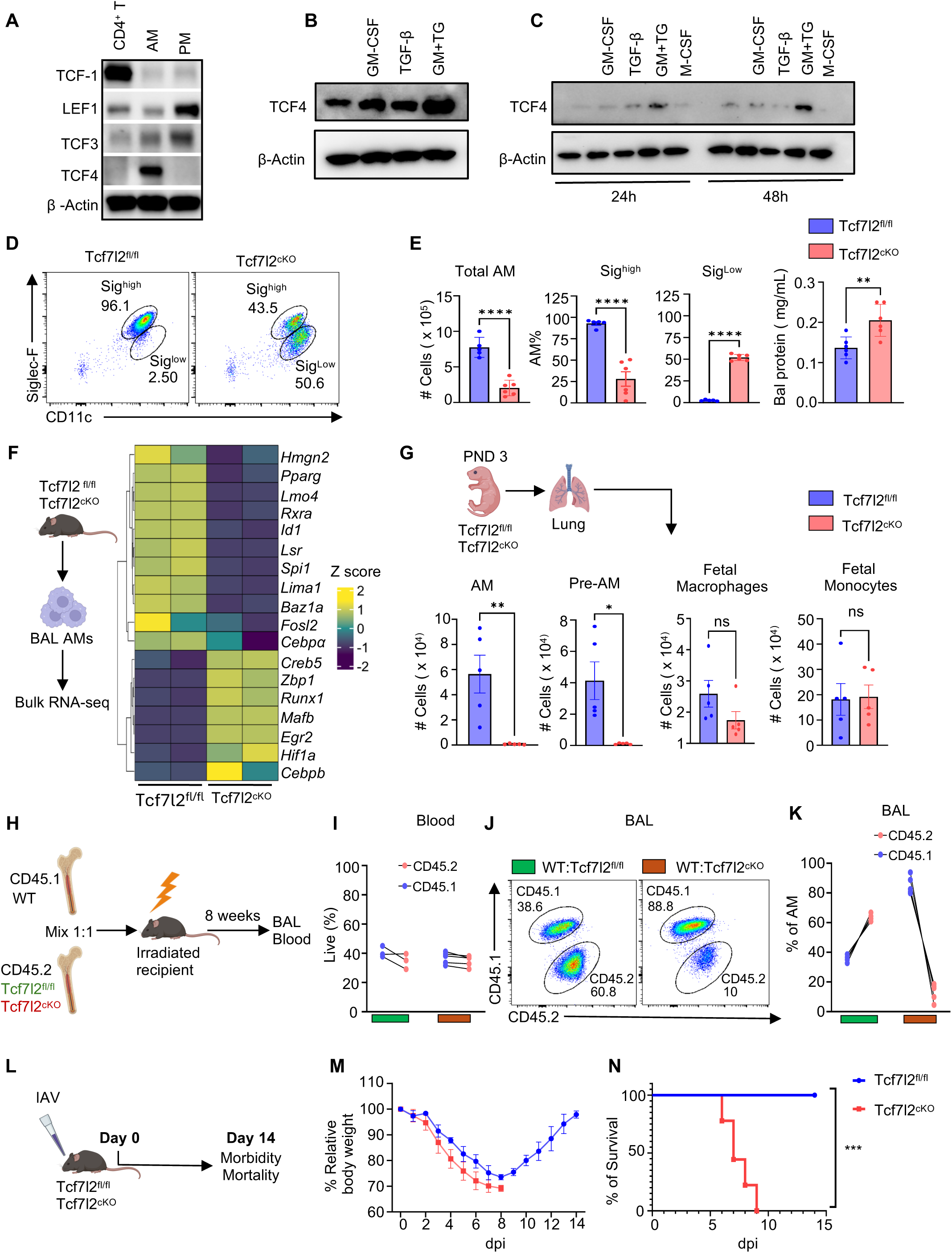
Intrinsic TCF4 is required for mature AM development and maturation. A. Representative western blot for TCF isoforms in primary mouse AMs, CD4 T cells, and peritoneal macrophages (PM). B. Representative western blot for TCF4 in primary mouse AMs without or with GM-CSF, TGF-β or both stimulation after 48h. C. Representative western blot for TCF4 in primary mouse monocytes without or with GM-CSF, TGF-β or both stimulation after 24h and 48h. D. Representative flow cytometry plots showing AM population in BAL cells isolated from Tcf7l2^fl/fl^ or Tcf7l2^cKO^ mice. E. Total AM numbers, Siglec F^high^, Siglec F^low^ AM, and total BAL protein were determined in naïve Tcf7l2^fl/fl^ (n=5) and Tcf7l2^cKO^(n=6) mice. F. Bulk RNA-seq analysis of AM isolated from Tcf7l2^fl/fl^ or Tcf7l2^cKO^ mice. A heatmap of k-means clustering of differentially expressed AM-associated transcription factors is shown. G. Schematic for lung immune cells isolation from PND3 mice. Down bar graph showing lung immune cells (AM, pre-AM, Fetal macrophages, and Fetal monocytes) from naïve PND3 Tcf7l2^fl/fl^ (n=5) or Tcf7l2^cKO^(n=5) mice. H. Schematic diagram for bone marrow (BM) chimera experiment (I-K). WT mice were lethally irradiated and reconstituted with CD45.1^+^ WT (50%) mixed with CD45.2^+^ Tcf7l2^fl/fl^ (50%) or CD45.2^+^ Tcf7l2^cKO^ (50%) BM cells. 8 weeks later, immune cell reconstitution in BAL fluid and blood was examined. I. Percentage of total frequency of CD45.1 and CD45.2 population from WT:Tcf7l2^fl/fl^ (n=3) and WT:Tcf7l2^cKO^ (n=5) mice in blood. J. Flow cytometry plot showing CD45.1 and CD45.2 population from WT:Tcf7l2^fl/fl^ and WT:Tcf7l2^cKO^ mice pre-gated on Siglec F^+^ and CD11c^+^ AM in BAL fluid. K. Percentage of CD45.1 and CD45.2 population from WT:Tcf7l2^fl/fl^ and WT:Tcf7l2^cKO^ mice pre-gated on Siglec F^+^ and CD11c^+^ AM in BAL fluid. L. Schematic for IAV infection model in Tcf7l2^fl/fl^ and Tcf7l2^cKO^ mice. M. Relative original body weight in percentages of Tcf7l2^fl/fl^ (n=5) or Tcf7l2^cKO^(n=9) mice post infection. N. Survival plot in Tcf7l2^fl/fl^ (n=5) or Tcf7l2^cKO^(n=9) mice after IAV infection. Data are representative of at least two independent experiments with similar results except F. *p*-values are represented as \**p*< 0.05, \*\**p*< 0.01, \*\*\**p*< 0.005, \*\*\*\**p*< 0.001.

To further study the role of TCF4, we checked the AM population in 8 weeks old mice in WT and myeloid-specific TCF4-deficient mice (Lyz2^cre^Tcf7l2^flox/flox^, Tcf7l2^cKO^). We found that in Tcf7l2^cKO^mice, there were defects in AM maturation and number, reflected by the decrease of Siglec F^high^ AMs, the emergence of Siglec F^low^ population and significantly lower AM number in bronchoalveolar lavage fluid (BAL) (Fig. 1D-E, S1I). In addition, the majority of AMs present in BAL of Tcf7l2^cKO^ mice were CD11b^+^, suggesting possible new recruitment of monocytes and/or failed maturation of monocytes into mature AMs in these mice (Fig. S1J). Also, Tcf7l2^cKO^ mice had accumulated protein content in BAL (Fig. 1E), further indicating defective AM functions in the absence of TCF4, as AMs are a primary cell type for clearing pulmonary surfactant proteins (*28*). Bulk RNAseq on these AMs showed TCF4-deficient AMs had defects in key AM transcription factors, including PPARγ (Fig. 1F), which is considered as the master transcription factor for AM development and maturation (*5*). On the other hand, HIF-1α, which is known to induce AM inflammatory programs (*11*), was upregulated in AMs isolated from Tcf7l2^cKO^ mice (Fig. 1F).

Next, we aimed to investigate the kinetics of AMs following TCF4 deficiency. AMs predominantly arise from fetal monocytes, transitioning through a pre-AM stage during the first week after birth (*8*). To investigate the role of TCF4 in early AM differentiation, we analyzed monocyte and macrophage populations in the lungs of Tcf7l2^fl/fl^ and Tcf7l2^cKO^ at postnatal day 3 (PND3). We found that the TCF4 deficiency led to defects in AMs (CD11c^+^ Siglec F^+^ Ly6C^−^ F4/80^+^) and pre-AM (CD11c^int^ Ly6C^−^ F4-80^+^ Siglec F^−^) population. However, both fetal monocyte (CD11c^−^ Ly6C^+^ Siglec F^−^ F4/80^−^) and fetal macrophages (CD11c^−^ Ly6C^+^ Siglec F^−^ F4/80^+^) showed non-significant changes in the lung (Fig. 1G). We further assessed AM numbers and phenotype during the juvenile stage (3 to 4 weeks postnatally). Our analysis revealed a significant reduction in both the frequency and numbers of mature AMs and the emergence of CD11b^+^ macrophages in Tcf7l2^cKO^ mice compared to WT controls (Fig. S1K-M). These data suggest that TCF4 is dispensable for fetal monocytes and macrophages seeding the lung, but critical for AM development and maturation.

We aimed to investigate whether TCF4 is inherently required for AM maturation. To this end, we performed bone marrow (BM) chimera in which BM cells were isolated from both CD45.1^+^ WT and Tcf7l2^cKO^ (CD45.2^+^) mice and injected in equal numbers into lethally irradiated CD45.2^+^ WT mice (Fig. 1H). Eight weeks after reconstitution, we observed comparable frequencies of CD45.1⁺ and CD45.2⁺ cells in the blood, confirming equal engraftment in recipient mice (Fig. 1I). However, we observed that mice received WT and Tcf7l2^fl/fl^ BM cells had almost equal AMs from WT donor and Tcf7l2^fl/fl^ mice (Fig 1J-K). In contrast, the majority of AMs were derived from WT origin (CD45.1) in mice that received BM cells from WT and Tcf7l2^cKO^ mice (Fig 1J-K). These data revealed that TCF4 is intrinsically required for mature AM development.

Mature AM presence is vital for the host’s resistance to influenza virus infection (*29*). To determine whether AM defects in Tcf7l2^cKO^ mice resulted in increased host susceptibility to influenza infection, we infected WT and Tcf7l2^cKO^ mice with sublethal influenza A/PR8/34 (H1N1) virus and checked for host morbidity and mortality post-infection until day 14 (Fig. 1L). Tcf7l2^cKO^ mice experienced greatly increased morbidity and mortality compared to those of control mice after IAV infection (Fig 1M-N). Together, our data discovered that TCF4 is critical for the development and maturation of AMs during homeostasis and host resistance to respiratory viral infection.

#### TCF4 is required for AM self-renewal

To further investigate the mechanisms by which TCF4 controls AM response, we performed bulk RNA sequencing of AM isolated from WT and Tcf7l2^cKO^ mice, both treated and untreated with Poly (I:C) (Fig. 2A). Our data found that Tcf7l2^cKO^ derived AMs exhibited inflammatory and hypoxia-related gene expression during homeostasis, which were further exacerbated after Poly (I:C) stimulation (Fig. S2A-B). Previously, we demonstrated that inflammatory phenotypes in AM negatively correlated with their proliferation capacity (*11*). Therefore, we examined the stemness and proliferative capacity of AM isolated from WT and Tcf7l2^cKO^ mice. Bulk RNAseq revealed an enrichment of embryonic stemness features in WT AMs compared to Tcf7l2^cKO^ AMs (Fig. 2B). *Mafb* and *c-Maf* have been previously shown to be expressed in fully differentiated macrophages to repress macrophage stemness (*30*). The expression of *Mafb* and *c-Maf* in Tcf7l2^cKO^ AMs was highly upregulated (Fig. 2C), suggesting these macrophages may have impaired self-renewal ability. Additionally, a colony-forming unit (CFU) assay showed that WT AMs could efficiently form colonies, whereas AMs isolated from Tcf7l2^cKO^ mice yielded very few colonies (Fig. 2D). Interestingly, AMs stimulated with GM-CSF and TGF-β had increased expression of both TCF4 and Ki67, and there was a strong co-expression of TCF4 and Ki67 in the nuclear region (Fig. S2C). Upon quantification, TCF4 expression was more enriched in Ki67^+^ AM (Fig. S2C). These data suggest that TCF4 expression may facilitate AM self-renewal. Consistent with the idea, we found that TCF4 deletion resulted in reduced proliferative capacity of Tcf7l2^cKO^ mice as compared to WT mice (Fig. 2E). Additionally, TCF4 deficiency caused decreased Ki67 expression with or without GM-CSF stimulation *in vitro* (Fig. S2D). These data together suggested that TCF4 is required for AM proliferation and TCF4 deficiency renders AM unresponsive to growth factor stimulation. Notably, GM-CSF, Siglec F, or TGF-β receptor genes were downregulated in Tcf7l2^cKO^ AMs (Fig. S2E), which may account for their lack of responsiveness to GM-CSF stimulation *in vitro*. In accordance, when BM cells were cultured under AM differentiation conditions including GM-CSF, TGF-β, and PPARγ agonist (*31*), TCF4-deficient BM cells failed to differentiate into Siglec F^high^ mature AM-like cells (Fig. S2F).

**Fig. 2.**
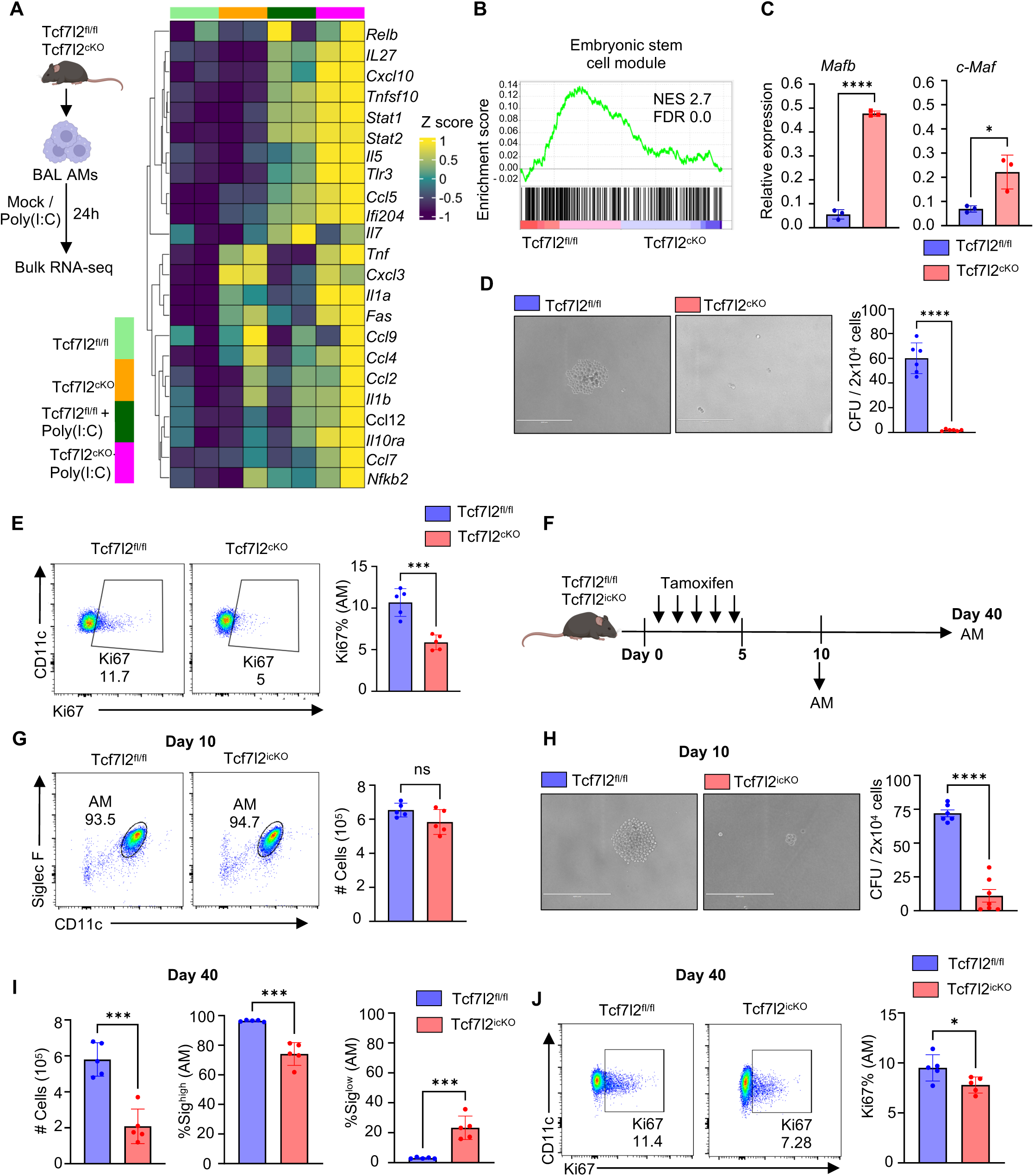
TCF4 is required for AM self-renewal. A. Schematic of Bulk RNA-seq analysis of Tcf7l2^fl/fl^ and Tcf7l2^cKO^ AMs. The Right Heatmap of k-means clustering of the differentially expressed top 100 genes is shown. B. Embryonic stem cell module of Tcf7l2^fl/fl^ and Tcf7l2^cKO^ AMs. C. *Mafb*/*c-Maf* gene expression in Tcf7l2^fl/fl^ and Tcf7l2^cKO^ AMs. D. Representative image of colony of Tcf7l2^fl/fl^ or Tcf7l2^cKO^ AMs. The right graph represents quantification of colony formation efficiency of Tcf7l2^fl/fl^ or Tcf7l2^cKO^ AMs. Scale bar 400 μm. E. Flow cytometry plot showing frequency of Ki67^+^ AM isolated from Tcf7l2^fl/fl^ or Tcf7l2^cKO^ mice. Right bar graph showing quantification of the frequency of Ki67^+^ AM F. Schematic of Tamoxifen administration (2mg/ml) in naïve Tcf7l2^fl/fl^ or Tcf7l2^icKO^ mice. G. Flow cytometry plot showing frequency of Siglec F^+^ CD11c^+^ AM in Tcf7l2^fl/fl^ (n=5) or Tcf7l2^icKO^ (n=5) mice BAL at day 10 post tamoxifen injection. Bar graph showing quantification of total cell number in Tcf7l2^fl/fl^ (n=5) or Tcf7l2^icKO^ (n=5) mice BAL. H. Representative image of CFU from AMs isolated from Tcf7l2^fl/fl^ or Tcf7l2^icKO^ mice at day 10 post tamoxifen injection. Right graph represents quantification of colony formation efficiency of AM isolated from Tcf7l2^fl/fl^ or Tcf7l2^icKO^ mice. Scale bar 400 μm. I. Bar graph showing quantification of total cell number, Siglec F^high^, and Siglec F^low^ frequency in Tcf7l2^fl/fl^ (n=5) and Tcf7l2^icKO^ (n=5) mice BAL fluid at day 40. J. Flow cytometry plot showing frequency of Ki67^+^ AM in Tcf7l2^fl/fl^ (n=5) or Tcf7l2^icKO^ (n=5) AM at day 40 post tamoxifen injection. Bar graph showing quantification of frequency of Ki67^+^ AM in Tcf7l2^fl/fl^ (n=5) or Tcf7l2^icKO^ (n=5) AMs. Data are representative of at least two independent experiments with similar results except A-B, *p*-values are represented as ns = non-significant, \**p*< 0.05, \*\*\**p*< 0.005,\*\*\*\**p*< 0.001.

It is possible that impaired AM development and reduced AM maturation in TCF4-deficient AMs may contribute to the diminished AM proliferation rather than the direct control of AM stemness by TCF4. To address this question, we generated inducible myeloid TCF4 deficiency (Tcf7l2^icKO^) by crossing Tcf7l2 floxed mice with Lyz2-Cre ERT2 mice. To this end, we deleted TCF4 in AMs of adult mice through the injection of Tamoxifen for 5 consecutive days, followed by 5-day rest (Fig. 2F). To validate TCF4 deletion in the inducible model, we isolated AMs from WT, Tcf7l2^cKO^, and tamoxifen-treated Tcf7l2^icKO^ mice and assessed TCF4 expression at day 10. AMs from both Tcf7l2^cKO^ and Tcf7l2^icKO^ mice exhibited a marked reduction in TCF4 expression compared to WT controls (Fig. S2G). Nevertheless, we observed no significant changes in AM numbers and the expression of CD11c and Siglec F in Tcf7l2^icKO^ mice as compared to WT mice (Fig. 2G), indicating that the acute depletion of TCF4 in mature AMs did not affect their short-term phenotypes. However, CFU assay on these AMs showed that TCF4 deficiency resulted in decreased CFU count, suggesting TCF4 deficiency causes defects in AM self-renewal (Fig. 2H). Consistently, AM numbers were significantly decreased in Tcf7l2^icKO^ mice 40 days following TCF4 deletion (Fig. 2I). Induced TCF4 deficiency resulted in the appearance of a subsets of AM population characterized by low expression of Siglec F, indicating that TCF4 deficiency resulted in gradual loss of AM maturation (Fig. 2I). In support of the idea, we found that AM isolated from Tcf7l2^icKO^ have less Ki67 staining, suggesting lesser proliferation capacity than the WT AM (Fig. 2J). Together, these data from inducible TCF4-deficient mice revealed a critical role of TCF4 in regulating AM proliferation and self-renewal.

#### TCF4 and **β**-catenin reciprocally regulate AM inflammation and self-renewal

The transcriptional responses induced by the Wnt/β-catenin pathway are primarily, if not exclusively, mediated by the TCF family members (*32*). Previously, we have found that β-catenin complexed with TCF4 in AMs under homeostatic conditions, however, under Wnt stimulation, β-catenin preferentially bound to HIF-1α instead (*11*). To further investigate the fate of TCF4 upon β-catenin deletion, we checked the expression of TCF4 upon β-catenin deletion in AMs. AMs isolated from myeloid-specific β-catenin-deficient mice (Ctnnb1^cKO^) mice were cultured in presence and absence of Poly (I:C) for 24h. Upon β-catenin deletion, AMs had increased expression of TCF4 (Fig. 3A). This suggested a negative regulation of β-catenin on TCF4 expression. In addition, we checked active β-catenin (non-phosphorylated form of β-catenin) expression and downstream HIF-1α (*33*) levels upon TCF4 deletion in AMs. We found that active β-catenin, and particularly its binding partner HIF-1α, were increased especially following Poly (I:C) stimulation in Tcf7l2^cKO^ AMs as compared to WT AMs (Fig 3B-D, Fig S3A). There was basal level significant increase in the HIF-1α levels in Tcf7l2^cKO^ AMs (Fig S3A), which further confirms our bulk-RNA seq (Fig. 1F). These data established that the reciprocal regulation of TCF4 and β-catenin-HIF-1α pathway in AMs, which is in contrast to the co-operative effects TCF4 and β-catenin in other cell types such as stem and cancerous cells.

**Fig. 3.**
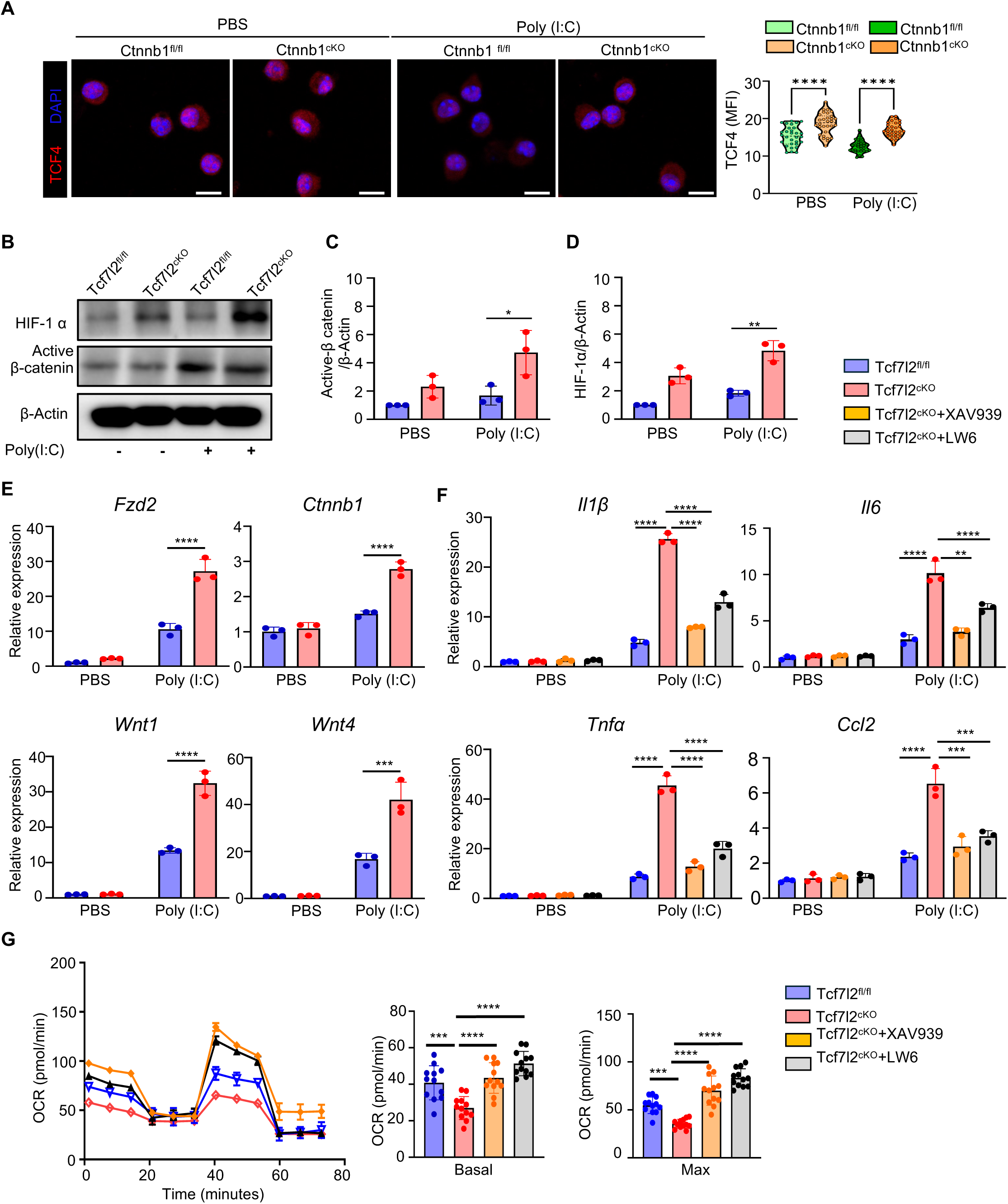
TCF4 and β-catenin reciprocally regulate AM inflammation and self-renewal. A. Representative IF images of TCF4 in Ctnnb1^fl/fl^ and Ctnnb^cKO^ AM stimulated with or without Poly(I:C). Violin plot representing TCF4 MFI in Ctnnb1^fl/fl^ or Ctnnb1^cKO^ AM stimulated with or without Poly(I:C). Scale bar 10 μm. B. Immunoblot of active β-catenin, HIF-1α and β-Actin in Tcf7l2^fl/fl^ or Tcf7l2^cKO^ AMs treated with or without Poly(I:C). C. Bar graph showing relative expression of active β-catenin in Tcf7l2^fl/fl^ or Tcf7l2^cKO^ AMs treated with or without Poly(I:C) quantified from three individual experiments. D. Bar graph showing relative expression of HIF-1α in Tcf7l2^fl/fl^ or Tcf7l2^cKO^ AMs. AMs treated with or without Poly(I:C) quantified from three individual experiments. E. Bar graph showing relative expression of Wnt signaling pathway genes (*Fzd2, Ctnnb1, Wnt1, Wnt4*) in Tcf7l2^fl/fl^ or Tcf7l2^cKO^ AMs treated with or without Poly(I:C). F. Bar graph showing relative expression of inflammatory cytokine expression (*Il1b, Il6, Tnf,* and *Ccl2*) in Tcf7l2^fl/fl^ or Tcf7l2^cKO^ AMs treated with or without β-catenin inhibitor (XAV939) or HIF-1α inhibitor (LW6). G. Graph showing oxygen consumption rate (OCR) measurements in WT and Tcf7l2^cKO^ AMs treated with or without β-catenin inhibitor (XAV939) or HIF-1α inhibitor (LW6). Right bar graph of OCR measurements in WT and Tcf7l2^cKO^ AMs treated with or without β-catenin inhibitor (XAV939) or HIF-1α inhibitor (LW6). Data are representative of at least two independent experiments with similar results, *p*-values are represented as \**p*< 0.05, \*\**p*< 0.01, \*\*\**p*< 0.005, \*\*\*\**p*< 0.001.

To examine the potential mechanisms underlying this reciprocal regulation of TCF4 and β-catenin function, we checked the expression of genes in Wnt pathway in Tcf7l2^cKO^ AM with or without Poly (I:C). We found that *Fzd2, Ctnnb1, Wnt1* and *Wnt4*, were upregulated in Tcf7l2^cKO^ AMs upon Poly (I:C) treatment (Fig. 3E). Thus, TCF4 could inhibit β-catenin activation likely through the regulation of β-catenin Wnt ligands (Wnt1 and Wnt4) and/or receptor (*Fzd2*) expression. To further confirm our understanding that Tcf7l2^cKO^ AMs have enriched in Wnt signaling pathway, we over-expressed TCF4 in WT AM, using adenovirus, and then checked the expression of these genes. We noticed that there was suppression in Wnt related genes pathway upon TCF4 overexpression (Fig. S3B-D).

We previously demonstrated that β-catenin and HIF-1α programs AM inflammatory activities (*11*). To this end, we utilized inhibitor of both β-catenin and HIF-1α, and investigated inflammatory cytokines expression, such as *Il1β*, *Il6*, *Tnfα and Ccl2* in TCF4-deficient AMs. TCF4 deletion in AMs had markedly increased inflammatory gene expression following Poly (I:C) treatment (Fig. 3F). However, β-catenin and HIF-1α inhibitor treatment in AMs dampened these cytokines gene expression, further suggesting that TCF4 represses β-catenin and HIF-1α activity to inhibit AM inflammation (Fig. 3F).

We next measured mitochondrial oxidative phosphorylation (OXPHOS) in WT and Tcf7l2^cKO^ AMs and detected a decrease in basal and maximum OCR (oxygen consumption rate). Notably, inhibition of β-catenin and HIF-1α led to elevated AM OCR in Tcf7l2^cKO^ AM following Poly(I:C) treatment (Fig. 3G, S3E), further confirming the roles of β-catenin and HIF-1α in mediating the effects of TCF4 depletion. Taken together, these data suggested that TCF4 antagonizes β-catenin function in AMs, thereby repressing AM inflammatory activity and promoting mitochondrial metabolism. Supporting this idea, we observed that β-catenin and TCF4 bound the same region of core glycolytic genes, such as *Enol* and *Hk2*, as identified by chromatin immunoprecipitation (ChIP) (Fig. S3F). Also, our ChIP assay showed that both β-catenin and TCF4 bound to key mitochondrial genes such as *Ndufs1* and *Ndufa4*, in the presence or absence of Wnt activator (Fig S3G). These data revealed that, rather than co-operating together downstream of the Wnt signaling as conventionally believed (*34*), TCF4 and β-catenin reciprocally regulate each other expression and function in AMs.

#### Acute respiratory viral infection suppresses AM TCF4 expression in mice and humans

AMs are critical for mitigating morbidity and mortality caused by acute respiratory viral infection (*29*). During infection, AM numbers as well as proliferation capacity are greatly reduced (*11*). As TCF4 regulates AM stemness features, we reasoned whether TCF4 expression is dysregulated in respiratory viral infections or not. In mouse models of IAV or SARS-CoV-2 (MA30 strain) infection, we observed that TCF4 expression in AMs was diminished at the peak of inflammation during acute infection (Fig. 4A-D). The markedly decreased TCF4 expression in AMs was observed on day 7 post-IAV infection when inflammation peaked (*35, 36*), followed by its restoration to basal levels by day 14, indicates a dynamic regulation of this transcription factor during infection (Fig. 4A-B). In SARS-CoV-2 infection, TCF4 expression showed a peak decrease on day 4, returning to close to basal levels by day 7 (Fig. 4C-D).

**Fig. 4.**
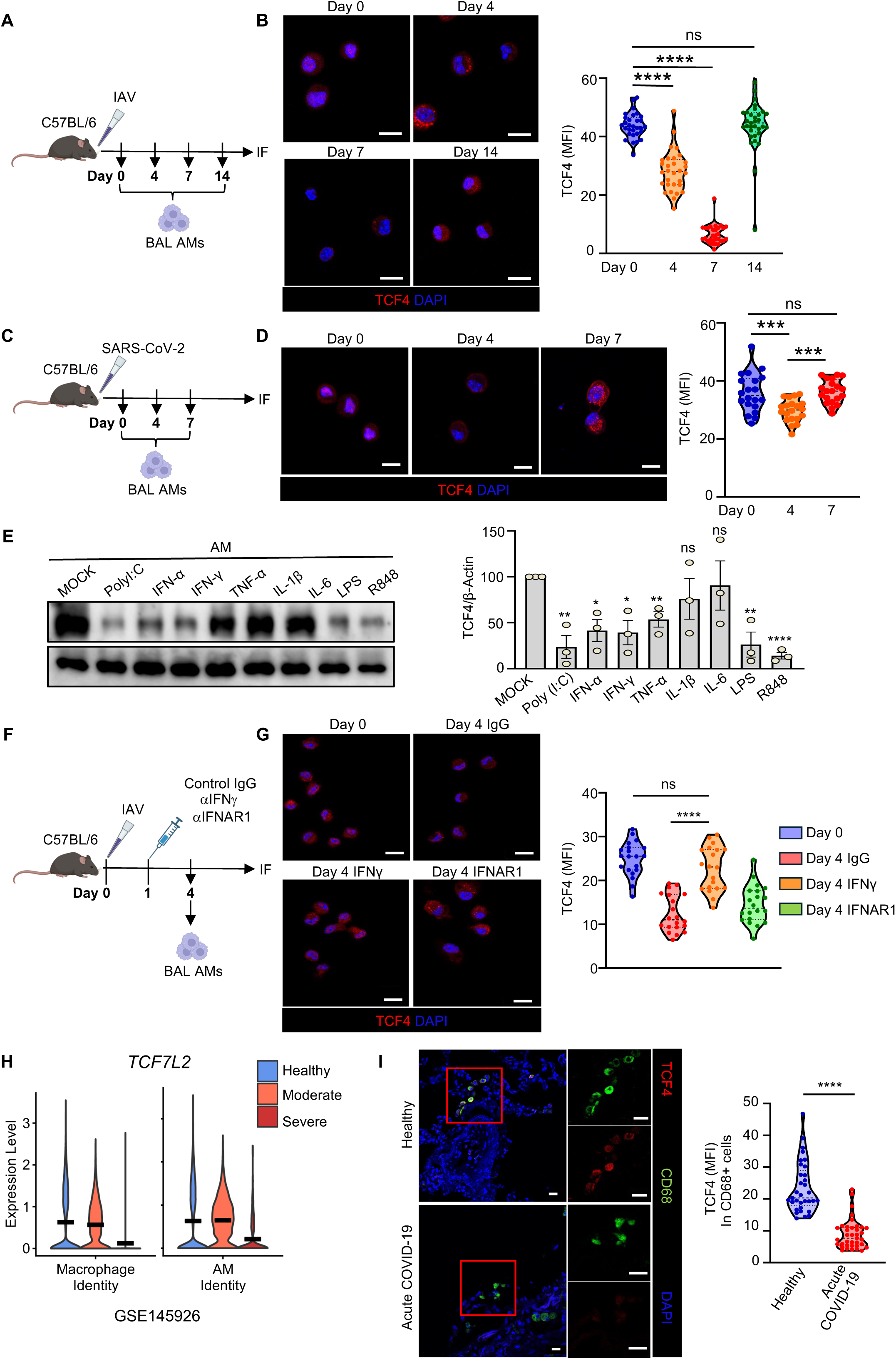
Severe viral infection causes AM TCF4 downregulation in mice and humans. A. Schematic diagram showing IAV infection kinetics with AM collection at day 0, 4, 7, and 14 for immunofluorescence (IF). B. Representative images of AM sorted at day 0, 4, 7 and 14 and stained with TCF4. Violin plots representing TCF4 MFI from AM. Scale bar 10 μm. C. Schematic diagram showing SARS-CoV-2 infection kinetics with AM collection at day 0, 4, and 7 for immunofluorescence. D. Representative images of AM sorted at day 0, 4, and 7 and stained with TCF4. Right violin plots representing TCF4 MFI. Scale bar 10 μm. E. Western blot showing TCF4 expression in WT AM stimulated with various TLR ligands and cytokines for 24h *in vitro*. Right bar graph showing TCF4 quantification in AM with respect to β-Actin in total three individual experiments. F. Schematic diagram showing IAV infection and antibody treatment with control IgG, anti-IFNAR1 or anti-IFNγ. AMs were sorted on day 4 and stained for TCF4. G. Representative images of AMs sorted at day 4 from IgG, anti-IFNγ, and anti-IFNAR1 groups and stained with TCF4. Scale bar 10 μm. Right violin plots representing TCF4 MFI from AM. H. Violin plot showing *TCF7L2* gene expression in lung total macrophages or AMs (right) from indicated COVID-19 BAL samples (GSE145926) (Healthy; healthy control; Moderate, moderate disease; Severe, severe disease). I. Representative confocal microscopy IF images of TCF4 staining in lung samples from non-COVID-19 control (n=3) and acute COVID-19 autopsies (n=5). Right Violin plots representing TCF4 MFI from CD68^+^ cells in control and acute COVID-19 autopsies. Scale bar 20 μm. Data are representative of at least two independent experiments with similar results. *p*-values are represented as ns = non-significant, \**p*< 0.05, \*\**p*< 0.01, \*\*\**p*< 0.005, \*\*\*\**p*< 0.001.

To investigate how viral infections diminish TCF4 expression in lung macrophages, we stimulated mouse AM *in vitro* with factors that model host signals induced by respiratory viral infections (Fig. 4E). We found that TLR ligands such as Poly (I:C), Lipopolysaccharide (LPS), and R848 and cytokines such as IFNα, IFNγ and TNF could significantly reduce the expression of TCF4 (Fig. 4E). As severe respiratory infection leads to outburst of IFN signaling, we further checked the role of these inflammatory cytokines in the downregulation of TCF4 expression in AMs *in vitro.* Type I or type II IFN markedly downregulated TCF4 expression in AMs when used alone or in combination *in vitro* in a dose-dependent manner (Fig S4A-B). Also, IFNγ resulted in enhanced HIF-1α and active β-catenin expression in Tcf7l2^cKO^ AMs, further suggestive reciprocal regulation of TCF4 and β-catenin in inflammatory conditions (Fig. S4C).

To determine whether IFNα or IFNγ mediates TCF4 downregulation during severe respiratory viral infection *in vivo,* we administered neutralizing anti-IFNγ or anti-IFNα/β receptor-blocking (anti-IFNAR1) antibodies following IAV infection. IFNγ neutralization resulted in increased TCF4 levels in AMs compared to the control group (Fig. 4F-G). IFNα/β receptor blockade was less effective than that of IFNγ neutralization (Fig. 4F-G). These findings indicate that excessive IFN responses, particularly the type II IFN, regulate TCF4 dynamics in AMs following respiratory viral infection. To determine the potential roles of IFNγ in regulating TCF4 expression in human AMs, we purified monocytes from a healthy donor and stimulated their differentiation into AM-like cells through the treatment of GM-CSF, TGF-β , and PPAR-γ agonist as previously reported (*18*) (Fig. S4D). Treatment of IFNγ in these AM-like cells also resulted in downregulation of TCF4 expression, indicating that IFNγ likely also inhibits TCF4 expression in human AMs (Fig. S4E).

Next, we examined whether severe respiratory viral infection in humans could result in diminished TCF4 expression in human AMs. To do so, we first utilized publicly available single-cell RNA sequencing (scRNA-seq) datasets (GSEA145926) to map the *TCF7L2* gene expression in lung total macrophages or AMs from BALs of healthy controls, COVID-19 individuals with moderate diseases, and COVID-19 individuals with severe diseases. Notably, *TCF7L2* gene expression was reduced in both total lung macrophages and the AM compartment in severe COVID-19 cases, but not in moderate cases, compared to healthy controls (Fig. 4H). To determine whether TCF4 expression is altered in macrophages during severe COVID-19, we performed immunofluorescence staining for TCF4 and CD68 in non-COVID-19 control lungs and acute COVID-19 autopsy lungs. CD68⁺ macrophages from COVID-19 autopsy lungs showed a marked reduction in TCF4 expression compared to controls (Fig. 4I). Together these data suggested that acute severe respiratory viral infection impeded AM TCF4 expression in mice and humans, potentially via exuberant IFN responses.

#### Inducible myeloid ablation of TCF4 impairs host recovery from respiratory viral infection

As Tcf7l2^cKO^ mice were highly susceptible to IAV infection due to impaired AM development, we next utilized Tcf7l2^icKO^ mouse model to examine whether acute ablation of TCF4 can lead to impaired host recovery from respiratory viral infection. To this end, control and Tcf7l2^icKO^ mice were injected with tamoxifen intraperitoneal (i.p) for 5 consecutive days and followed by 5 days rest (Fig. 5A). The Tcf7l2^icKO^ mice had a comparable number of AMs to control mice at this time point (Fig 2G). Next, to investigate the role of acute TCF4 deletion on host recovery upon respiratory viral infection, we infected WT and Tcf7l2^icKO^ mice with SARS-CoV-2 (Fig. 5A). In case of SARS-CoV-2 infection, Tcf7l2^icKO^ mice lost significantly more weight and exhibited increased mortality (Fig. 5B). Furthermore, analysis of the BAL at 10 days post infection (d.p.i) revealed a decreased frequency of AMs in the BAL (Fig 5C). Consistently, Tcf7l2^icKO^ mice showed extensive lung inflammation as compared to WT counterparts (Fig. 5D). Upon further analysis, there were less Siglec F^high^ frequency and cells count in Tcf7l2^icKO^ infected mice. On the contrary, there were more Siglec F^low^ AM frequency and number, which was accompanied by an increased trend in Ly6C^high^ monocytes (Fig. S5A).

**Fig. 5.**
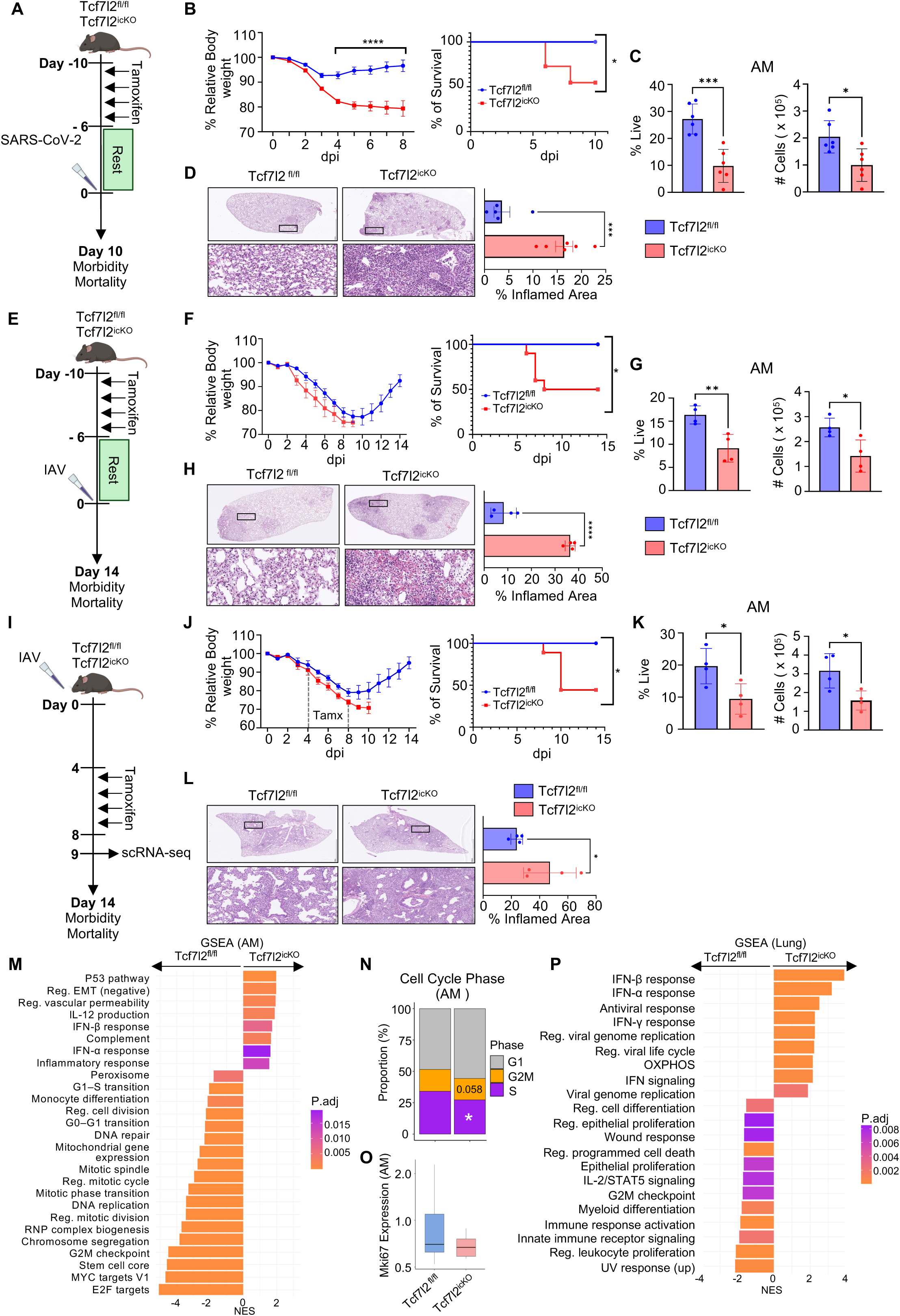
Inducible TCF4 ablation impairs host recovery from respiratory viral infection. A. Schematic of SARS-CoV-2 infection model in Tcf7l2^fl/fl^ or Tcf7l2^icKO^ mice. The mice were injected with tamoxifen for 5 consecutive days, followed by 5 days of rest. The mice were subsequently infected with SARS-CoV-2 and checked for morbidity and mortality until 10 d.p.i. B. Left, relative percentages of body weight loss in Tcf7l2^fl/fl^ (n=14) or Tcf7l2^icKO^ (n=15) mice. Right, survival plot of Tcf7l2^fl/fl^ (n=11) or Tcf7l2^icKO^ (n=11) mice after SARS-CoV-2 infection. C. Bar graph showing quantification of frequency and total AM number in Tcf7l2^fl/fl^ (n=6) and Tcf7l2^icKO^ (n=6) mice BAL fluid on day 10. D. Representative image for H&E section from Tcf7l2^fl/fl^ or Tcf7l2^icKO^ at 10 d.p.i. Bar graph on right showing % disrupted area in Tcf7l2^fl/fl^ (n=5) or Tcf7l2^icKO^ (n=6) E. Schematic of IAV infection model in Tcf7l2^fl/fl^ or Tcf7l2^icKO^ mice. The mice were injected with tamoxifen for 5 consecutive days, followed by 5 days of rest. The mice were subsequently infected with IAV and checked for morbidity and mortality. F. Left, relative body weight loss in percentages in Tcf7l2^fl/fl^ (n=9) or Tcf7l2^icKO^ (n=10) mice. Right, survival plot in Tcf7l2^fl/fl^ (n=9) or Tcf7l2^icKO^ (n=10) mice after IAV infection. G. Bar graph showing quantification of frequency and total AM number in Tcf7l2^fl/fl^ (n=4) or Tcf7l2^icKO^ (n=4) mice BAL on day 14. H. Representative image for H&E section from Tcf7l2^fl/fl^ or Tcf7l2^icKO^ mice at 14 d.p.i. Bar graph on the right showing % disrupted area in Tcf7l2^fl/fl^ (n=4) or Tcf7l2^icKO^ (n=4). I. Schematic of acute deletion of Tcf7l2 in the IAV infection model. The mice were infected with IAV (150 PFU) on day 0 and injected with tamoxifen for 5 consecutive days on day 4 and checked for morbidity and mortality until 14 d.p.i. A group of mice was sacrificed on day 9 for single-cell RNA sequencing (scRNA). J. Left, relative body weight loss in percentages in Tcf7l2^fl/fl^ (n=8) or Tcf7l2^icKO^ (n=9) mice. Right, survival plot in Tcf7l2^fl/fl^ (n=8) or Tcf7l2^icKO^ (n=9) mice after IAV infection. K. Bar graph showing quantification of frequency and total AM number in Tcf7l2^fl/fl^ (n=4) or Tcf7l2^icKO^ (n=4) mice BAL fluid at 14 d.p.i. L. Representative image for H&E section from Tcf7l2^fl/fl^ or Tcf7l2^icKO^. Bar graph on right showing % disrupted area in Tcf7l2^fl/fl^ (n=4) or Tcf7l2^icKO^ (n=4). M-P. Lung scRNAseq analysis was performed at 9 d.p.i. M. GSEA showing upregulated pathways in AM from IAV infected Tcf7l2^fl/fl^ or Tcf7l2^icKO^ mice. N. Bar graph showing proportion of cell cycle phase in AM from IAV infected Tcf7l2^fl/fl^ or Tcf7l2^icKO^ mice. O. Bar graph showing expression of Mki67 in AM from IAV infected Tcf7l2^fl/fl^ or Tcf7l2^icKO^ mice. P. GSEA showing upregulated pathways in total lung cells from IAV infected Tcf7l2^fl/fl^ or Tcf7l2^icKO^ mice. Data are representative of at least two independent experiments with similar results except M-P, *p*-values are represented as \**p*< 0.05, \*\*\**p*< 0.005, \*\*\*\**p*< 0.001.

Similarly, WT and Tcf7l2^icKO^ mice received i.p tamoxifen injections for five consecutive days, followed by a five-day rest period prior to IAV infection (Fig. 5E). Tcf7l2^icKO^ mice infected with IAV also showed more weight loss as compared to WT mice, and the majority of Tcf7l2^icKO^ mice succumbed to the infection post day 9 (Fig. 5F). Similarly, Tcf7l2^icKO^ mice showed decreased AM frequency count and numbers (Fig. 5G), and a trend in the increased population of Ly6C^low^ monocytes (Fig. S5B). Furthermore, the lungs of Tcf7l2^icKO^ mice showed significantly greater inflammation and damage compared to their WT counterparts (Fig. 5H).

We also started tamoxifen treatment from day 4 to day 8 and monitored the recovery of these mice (Fig. 5I). Again, Tcf7l2^icKO^ mice failed to recover from the infection in this setting, and Tcf7l2^icKO^ mice observed a lower frequency and number of AMs as compared to WT mice at day 14 (Fig. 5J-K). In addition, this time we observed an increased number of Ly6C^high^ monocytes (Fig. S5C). The H&E section from the lungs of Tcf7l2^icKO^ mice also showed extensive inflammation as compared to the WT mice (Fig. 5L). Taken together, these data established that continued TCF4 expression is critical for the maintenance of the AM compartment *in vivo* and host recovery from respiratory viral infection.

Because Tcf7l2^icKO^ mice failed to recover following influenza infection, we next sought to define the transcriptional programs underlying this phenotype using single cell RNA sequencing (scRNAseq) (Fig. 5I). As most Tcf7l2^icKO^ mice succumbed after day 9 post-infection (Fig. 5J), we performed lung scRNAseq at this time point and checked for cell distribution in WT and Tcf7l2^icKO^ (Fig. S5D–E). Gene set enrichment analysis (GSEA) revealed that AMs from Tcf7l2^icKO^ mice were enriched for p53 pathway genes, consistent with cell-cycle arrest and senescence (*37*), as well as inflammatory signatures characterized by complement activation, IL-12 production, and type I interferon responses (Fig. 5M). By contrast, AMs from Tcf7l2^fl/fl^ mice exhibited enrichment of active cell-cycle programs, stem cell core transcriptional modules, and DNA repair pathways (Fig. 5M). Notably, AMs from Tcf7l2^fl/fl^ mice showed significant enrichment in the S phase, required for DNA replication, and expressed the proliferation marker Mki67 (Fig. 5N-O). These cells also retained expression of canonical AM identity genes (*Itgax*, *Adgre1*, *Cd68*) and upregulated *Plet1*, a secreted factor that promotes lung repair (*38*) (Fig. S5F). In contrast, volcano plot analysis identified a subset of top DEGs in Tcf7l2^icKO^ AMs associated with inflammasome activation and interferon-stimulated gene (ISG) responses (Fig. S5G). Together, these data suggest that TCF4 is required for AM proliferative and reparative responses after viral infection, while repressing AM inflammatory activities that lead to enhanced pulmonary injury after infection.

Consistently, at the whole-lung level, GSEA confirmed broad enrichment of type I and type II interferon responses in Tcf7l2^icKO^ mice, whereas lungs from Tcf7l2^fl/fl^ controls displayed enrichment in epithelial proliferation, wound-healing programs, and myeloid differentiation pathways (Fig. 5P). Together, these data suggest that TCF4 expression in AMs engages proliferative and reparative pathways that promote tissue recovery after viral injury.

#### Myeloid TCF4 limits lung PASC development

As introduced above, the current understanding of the mechanisms leading to the development of chronic lung diseases after acute viral infection remains largely undefined. Our prior studies have revealed a critical role of AM in lung repair and modulating the development of chronic lung diseases (*9, 39*). We thus hypothesize that impaired TCF4 expression and/or function in macrophages may lead to the development of chronic lung conditions after acute infection. To this end, we infected tamoxifen-treated WT control and Tcf7l2^icKO^ mice with SARS-CoV-2 or IAV, and studied post viral lung sequelae at 21 d.p.i. or 42 d.p.i. respectively as we have previously reported (*18, 40*). Mice that survived acute infection were sacrificed on 21 d.p.i or 42 d.p.i respectively to examine Siglec F^high^ and Siglec F^low^ population in AMs (Fig. 6A-F). AMs from Tcf7l2^icKO^ mice were mainly Siglec F^low^ AMs, suggesting the impaired repopulation of mature AMs or failure of AM maturation from recruited monocytes in the absence of TCF4 (Fig. 6B-C). Similar findings were observed post-acute IAV infection (Fig. 6E-F).

**Fig. 6.**
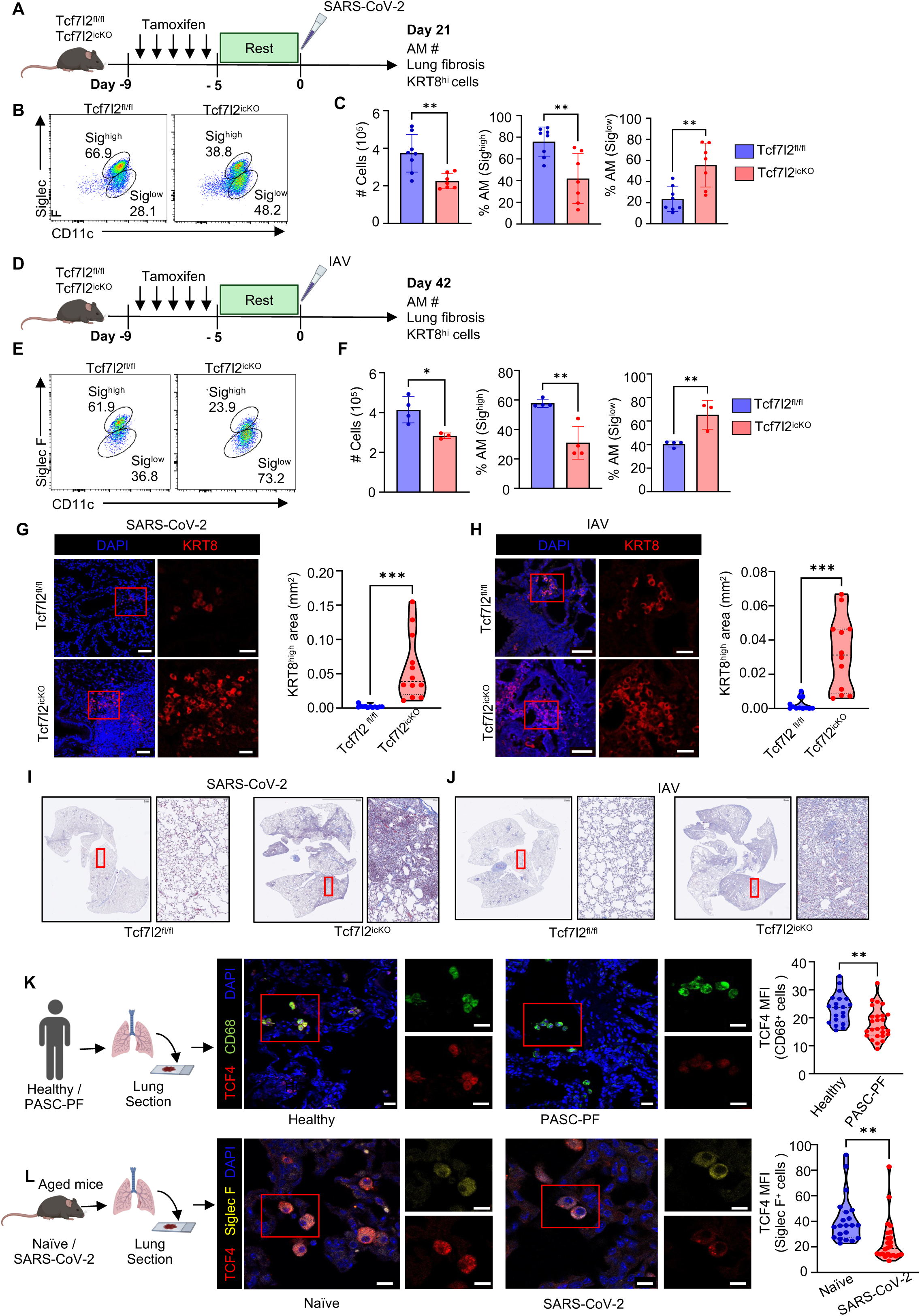
Myeloid TCF4 limits post viral lung fibrosis, and human PASC-PF patients exhibit persistent impairment of macrophage TCF4 expression. A. Schematic of long-term SARS-CoV-2 infection model in Tcf7l2^fl/fl^ or Tcf7l2^icKO^ mice. The mice were injected with tamoxifen (2mg/mL) i.p for 5 consecutive days followed by 5 days of rest. The mice were subsequently infected with SARS-CoV-2 and checked for immune cells and lung pathology at 21 d.p.i. B. Flow cytometry plot showing frequency of Siglec F^+^ CD11c^+^ AM in Tcf7l2^fl/fl^ (n=8) or Tcf7l2^icKO^ (n=7) mice BAL at 21 d.p.i. C. Bar graph showing quantification of AM # and frequency of Siglec F^high^ or Siglec F^low^ AM in Tcf7l2^fl/fl^ (n=8) or Tcf7l2^icKO^ (n=7) mice BAL. D. Schematic of long-term IAV infection model in Tcf7l2^fl/fl^ or Tcf7l2^icKO^ mice. The mice were injected with tamoxifen (2mg/mL) i.p for 5 consecutive days, followed by 5 days of rest. The mice were then infected with IAV and checked for immune cells and lung pathology on 42 d.p.i. E. Flow cytometry plot showing frequency of Siglec F^+^ and CD11c^+^ AM in Tcf7l2^fl/fl^ (n=4) and Tcf7l2^icKO^ (n=3) mice BAL fluid at 42 d.p.i. F. Bar graph showing quantification of AM # and frequency of Siglec F^high^ or Siglec F^low^ AM in Tcf7l2^fl/fl^ (n=4) or Tcf7l2^icKO^ (n=3) mice BAL. G. Representative confocal microscopy IF images of KRT8 staining in lung samples from SARS-CoV-2 infected Tcf7l2^fl/fl^ (n=4) or Tcf7l2^icKO^ (n=4) mice at 21 d.p.i. Violin plot showing quantification of KRT8 staining in lung samples. Scale bar 50 μm or 20 μm. H. Representative confocal microscopy IF images of KRT8 staining in lung samples from IAV-infected Tcf7l2^fl/fl^ (n=4) or Tcf7l2^icKO^ (n=3) mice at 42 d.p.i. Scale bar 50 μm or 20 μm. Violin plot showing quantification of KRT8 staining in lung samples. I. Representative Masson Trichome staining indicating lung fibrosis in Tcf7l2^fl/fl^ or Tcf7l2^icKO^ mice after SARS-CoV-2 infection at 21 d.p.i. J. Representative Masson Trichome staining indicating lung fibrosis in Tcf7l2^fl/fl^ or Tcf7l2^icKO^ mice after IAV infection at 42 d.p.i. K. Representative confocal IF images of CD68 and TCF4 staining in lung samples from non-COVID-19 control (HC) (n=3) or PACS-PF patients (n=5). Violin plot showing quantification of TCF4 MFI in CD68^+^ cells. Scale bar 20 μm. L. Representative confocal IF images of Siglec F and TCF4 staining in lung samples from naïve and 21 d.p.i SARS-CoV-2 infected mice. Scale bar 10 μm Violin plot showing quantification of TCF4 MFI in Siglec F^+^ cells. Data are representative of at least two independent experiments with similar results, *p*-values are represented as \**p*< 0.05, \*\**p*< 0.01, \*\*\**p*< 0.005.

We recently found that the persistence of transitional epithelial cells is the hallmark of post-viral lung sequelae in mice and humans (*19*). In consistency, IF of lung section of Tcf7l2^icKO^ mice showed a markedly increased accumulation of dysplastic KRT8^high^ transitional cells in the tissue of Tcf7l2^icKO^ infected mice compared to those of WT mice post SARS-CoV-2 or IAV infection (Fig. 6G-H). Our findings indicate that loss of Tcf7l2 arrests AM maturation, leading to the buildup of immature populations that fail to complete differentiation. This defective trajectory fuels the emergence of pathogenic KRT8⁺ transitional cells, thereby predisposing the lung to fibrotic remodeling. Consistently with the notion, Masson trichrome staining of these lung sections revealed increased lung fibrosis in the Tcf7l2^icKO^ mice post-acute infection (Fig. 6I-J). Furthermore, enhanced vascular leakage and the presence of chronic lung inflammation and injury were observed in Tcf7l2^icKO^ mice as compared to WT mice post-acute SARS-CoV-2 or IAV infection (Fig. S6A-D).

#### PASC-PF patients and PASC-PF mice exhibit persistent impairment of macrophage TCF4 expression

As acute COVID-19 patients exhibit decreased expression of TCF4 at the mRNA and protein level (Fig. 4H-I), we were curious to know if PASC patients with extensive pulmonary fibrosis (PASC-PF) (*41*), still had defects in the macrophage expression of TCF4. Analysis of publicly available dataset (GSE224955) from the lungs of PASC-PF patients suggested a decrease in expression of *TCF7L2* in lung macrophages of PASC-PF patients (Fig. S6E). To further confirm this, we performed IF imaging on macrophage TCF4 expression in lung sections from these PASC-PF patients and observed that TCF4 expression was still compromised in CD68^+^ macrophages of those patients compared to healthy controls (Fig. 6K). We recently reported that aged mice infected with SARS-CoV-2 developed notable post viral lung fibrosis at 3 weeks post infection, which represents a PASC-PF mouse model (*18*). Therefore, we utilized this mouse model and examined TCF4 expression in AMs. Similar to what was observed in humans, we found that Siglec F^+^ AMs from mice with chronic lung sequelae exhibited persistent reduced TCF4 expression compared to those of naïve mice (Fig. 6L, S6F, G). Taken together, these data revealed that proper macrophage TCF4 expression is likely important for preventing the development of chronic lung sequelae after acute COVID-19 in mice and humans.

#### TCF4 overexpression promotes AM self-renewal and mitigates severe viral infection

We have shown how TCF4 and Ki67 expressions were tightly co-regulated, and AMs lost their ability of self-renewal upon TCF4 deletion. Next, we sought to determine whether enforced TCF4 expression could facilitate AM self-renewal and inhibit severe host disease development after acute respiratory viral infection. To do this, we transduced WT AMs with control GFP-expressing adenovirus or TCF4-expressing adenovirus (Adv-Tcf7l2). Adenoviral delivery was effective *in vivo* (Fig. S7A), and Adv-Tcf7l2–transduced AMs maintained elevated TCF4 expression even in the absence of GM-CSF and under Poly(I:C) challenge (Fig. S7B).

Next, we checked the proliferation ability of AMs transduced with control or Adv-Tcf7l2 AM *in vitro* and observed a significant growth of cell number with Adv-Tcf7l2 (Fig. S7C-D). In accordance with the above results, we observed increased Ki67 expression in bulk AMs that were cultured for 48h (Fig S7E). As in this experiment, there were some non-transduced GFP^−^ AMs, we then sorted transduced GFP^+^ AMs and again examined Ki67 expression (Fig. S7F). Adv-Tcf7l2 transduction markedly enhanced TCF4 and Ki67 expression, with prominent colocalization of TCF4 and Ki67 observed in sorted GFP⁺ AMs (Fig. 7A). When cultured in the presence or absence of GM-CSF, Tcf7l2-transduced AMs exhibited elevated Ki67 staining *in vitro* (Fig. S7G). To further assess proliferative capacity, we performed CFU assays and found that enforced TCF4 expression significantly augmented both the number and size of AM-derived colonies (Fig. 7B). Next, we checked if TCF4 over-expression can also dampen the inflammatory response in AMs upon stimulation with Poly (I:C). Again, for this study, we utilized sorted GFP^+^ AMs and treated them with or without Poly (I:C). Following TCF4 over-expression, the expression of those inflammatory genes (*Il1β*, *Tnf, Il6*, *Ccl2*) were significantly downregulated (Fig. 7C). These data collectively suggested that enforced TCF4 expression can facilitate AM replication and self-renewal, while inhibiting their inflammatory activities.

**Fig. 7.**
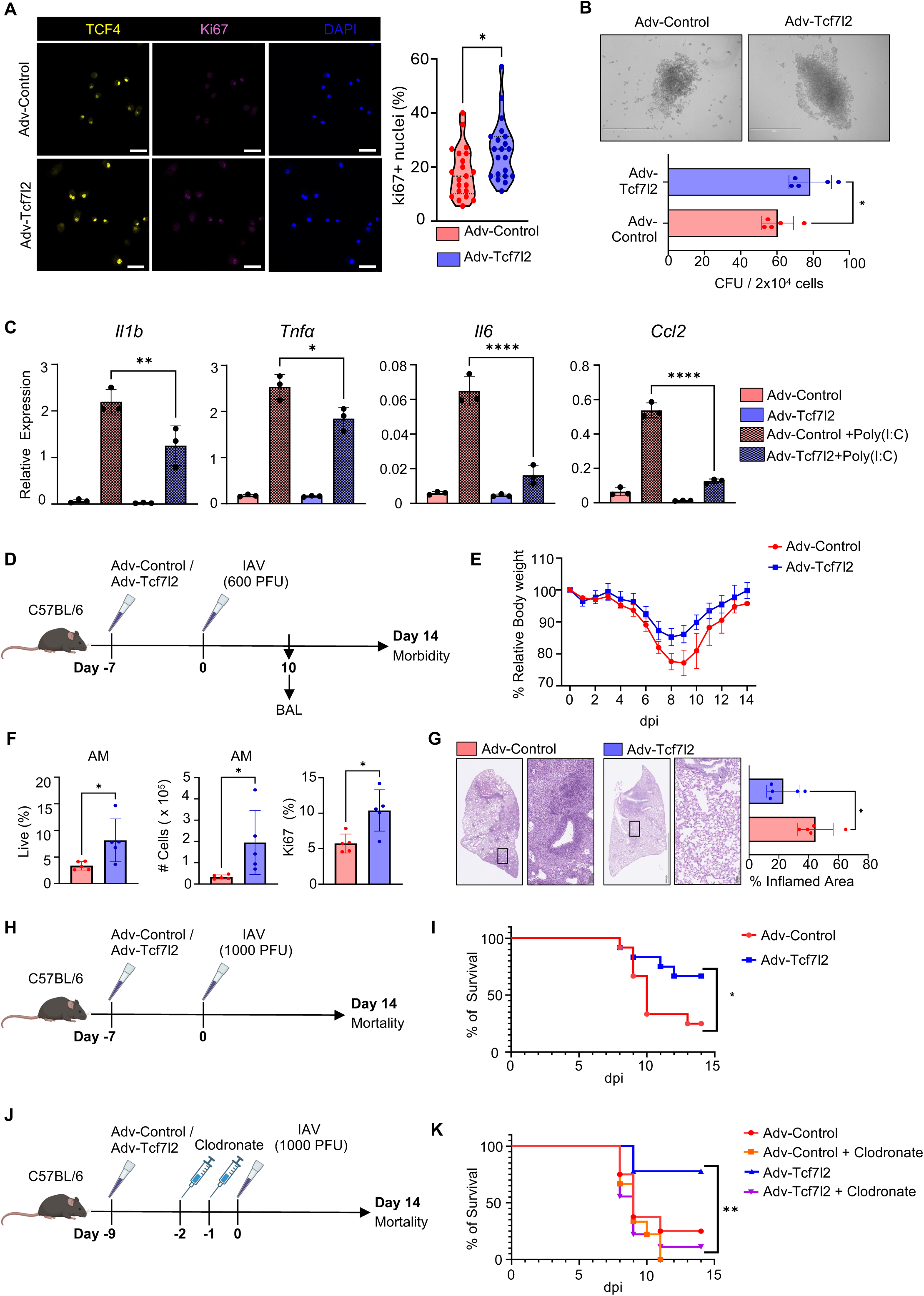
TCF4 overexpression promotes AM self-renewal and accumulation, and inhibits severe viral infection. A. Representative confocal microscopy IF images of GFP^+^ AMs sorted from WT mice transduced with Adv-Control or Adv-Tcf7l2 and stained with TCF4 and Ki67. Violin plot showing Ki67^+^ AM in transduced GFP^+^ AM. Scale bar 10 μm. B. Representative image of colony from GFP^+^ AMs sorted from WT mice transduced with Adv-Control or Adv-Tcf7l2. Right bar graph showing the number of colonies per 20,000 sorted GFP^+^ AM. Scale bar 1000 μm. C. Inflammatory cytokine gene expression (*Il1b*, *Tnf*, *Il6* and *Ccl2*) in transduced GFP^+^ AMs sorted from Adv-Control or Adv-Tcf7l2 infected mice. AMs were stimulated with or without Poly(I:C) for 24h *in vitro*. D. Schematic model showing WT mice transduced with Adv-Control or Adv-Tcf7l2 and then infected with IAV (600PFU). Mice morbidity was examined 14 days post IAV infection. E. Relative percentage of body weight loss of WT mice transduced with Adv-Control (n=4) or Adv-Tcf7l2 (n=5) and then infected with IAV (600PFU). F. Bar graph showing quantification of frequency, total AM number, and Ki67^+^ AM frequency in BAL of WT mice transduced with Adv-Control (n=5) or Adv-Tcf7l2 (n=5) and subsequently infected with IAV (600PFU) on day 10 post IAV infection. G. Representative image for H&E section from WT mice transduced with Adv-Control or Adv-Tcf7l2 and then infected with IAV (600PFU). Bar graph on the right showing % disrupted area. H. Schematic model showing WT mice transduced with Adv-Control or Adv-Tcf7l2 and then infected with IAV (1000PFU). Mice mortality was examined until day 14. I. Survival plot in WT mice transduced Adv-Control (n=12) or Adv-Tcf7l2 (n=12) and then infected with IAV (1000PFU). J. Schematic model showing AM depletion using intranasal clodronate. Mice were transduced with Adv-Control or Adv-Tcf7l2 followed by intranasal clodronate depletion. Mice were then infected with IAV (1000PFU) and mortality was examined until day 14 post IAV infection. K. Survival plot in WT mice transduced with Adv-Control alone (n=8), Adv-Tcf7l2 alone (n=9), Adv-control with intranasal clodronate (n=9), or Adv-Tcf7l2 with intranasal clodronate delivery (n=9). Data are representative of at least two independent experiments with similar results, *p*-values are represented as \**p*< 0.05, \*\**p*< 0.01, \*\*\*\**p*< 0.001.

As severe viral infection impairs AM stemness and promotes lung injury, we next sought to examine whether the overexpression of TCF4 could ameliorate virus-induced lung injury following respiratory viral infection. In this study, WT mice were transduced with Adv-control or Adv-Tcf7l2, and then infected with a high dose of IAV (600 PFU) (Fig. 7D). The majority of mice transduced with Adv-Tcf7l2 showed a remarkable resistance to weight loss (Fig. 7E). To further assess whether TCF4 overexpression can rescue AMs following IAV infection, we analyzed AM frequency and number at 10 d.p.i. Overexpression of TCF4 *via* Adv-Tcf7l2 resulted in increased AM frequency and total AM number, along with a higher proportion of Ki67⁺ AMs in the BAL, indicating enhanced proliferation *in vivo* (Fig. 7F). Lungs isolated from these mice demonstrated reduced inflammation as observed in H&E section (Fig. 7G). We also utilized a lethal dose (1000 PFU) on these transduced mice to examine if Adv-Tcf7l2 can improve host mortality after IAV infection (Fig. 7H). Mice transduced with Adv-Tcf7l2 were able to recover after acute infection, while the majority of mice with Adv-control transduction succumbed to the infection (Fig. 7I, S7H). To determine whether this rescue of mice from severe respiratory infection was AM-dependent, we depleted AMs after the adenovirus transduction with clodronate treatment. AM depletion in Adv-Tcf7l2 mice failed to provide protection in mice, suggesting the rescue of host mortality by Adv-Tcf7l2 transduction is dependent on the presence of AMs (Fig. 7K, S7I).

#### Targeting TCF4 to dampen age-associated chronic lung diseases after infection

We have previously shown that AMs from aged mice demonstrated impaired self-renewal capacity and diminished responsiveness to GM-CSF (*42*). Interestingly, TCF4 expression was impaired in aged AMs as compared to young AMs (Fig. 8A). To determine whether the provision of TCF4 in aged AMs could facilitate their proliferation, we infected aged AMs with Adv-control or Adv-Tcf7l2 and found that TCF4 over-expression in GFP+ AM (Fig. S8A). We next assessed the proliferative capacity of young and aged AMs transduced with either Adv-Control or Adv-Tcf7l2 *in vitro*. Aged AMs transduced with Adv-Tcf7l2 exhibited a significant increase in cell number compared with Adv-Control (Fig. S8B–C). Consistently, Ki67 expression was elevated in bulk AM cultures at day 4 (Fig. S8D).

**Fig. 8.**
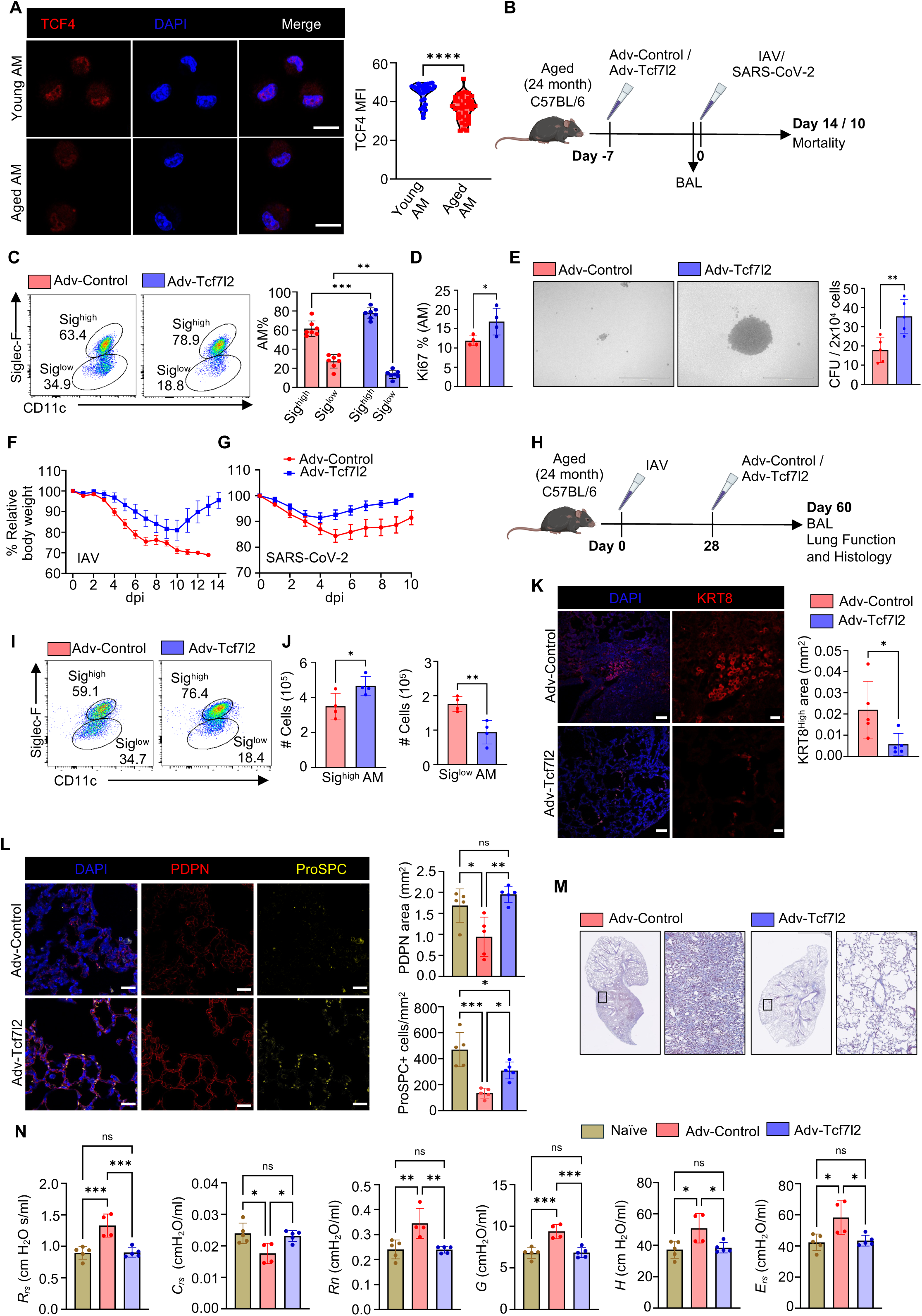
Targeting TCF4 to enhance AM self-renewal and limit age-associated acute and chronic lung diseases after infection. A. Representative confocal microscopy IF images of AM isolated from young and aged mice stained with TCF4. Violin plot showing quantification of TCF4 MFI. B. Schematic model showing aged WT mice transduced with Adv-Control or Adv-Tcf7l2. Each group of mice was sacrificed on day 0, and others were infected with IAV (150PFU) or SARS-CoV-2 (1000 PFU) on day 0. Mice morbidity was examined until day 14 or 10, respectively. C. Flow cytometry plot showing frequency of Siglec F^+^ and CD11c^+^ AM in naïve aged mice transduced with Adv-Control (n=7) and Adv-Tcf7l2 (n=7) mice. Bar graph showing frequency of Siglec F^high^ and Siglec F^low^ AMs in aged mice treated with Adv-Control (n=7) and Adv-Tcf7l2 (n=7) mice. D. Bar graph showing frequency of Ki67^+^ AM from sorted GFP^+^ AM isolated from WT aged mice transduced with Adv-Control (n=4) and Adv-Tcf7l2 (n=4). E. Representative image of CFU in GFP^+^ AMs sorted from WT mice transduced with Adv-Control or Adv-Tcf7l2. Right bar graph showing the number of CFU per 20,000 sorted GFP^+^ AM. F. Relative body weight loss in percentage in Adv-Control (n=10) or Adv-Tcf7l2 (n=10) infection in aged WT mice followed by IAV infection (150 PFU) on day 0. G. Relative percentage of body weight loss in Adv-Control (n=6) or Adv-Tcf7l2 (n=6) treated aged mice followed by SARS-CoV-2 infection (1000 PFU) on day 0. H. Schematic diagram for chronic model of IAV infection. Aged WT mice were infected on day 0, following recovery mice were infected with Adv-Control or Adv-Tcf7l2 day 28. Lung function was assessed, followed by BAL fluid collection and lung isolation on day 60. I. Flow cytometry plot showing frequency of Siglec F^+^ and CD11c^+^ AM isolated on day 60 from IAV-infected aged mice reinfected with Adv-Control (n=4) or Adv-Tcf7l2 (n=4). J. Bar graph showing frequency of Siglec F^high^ and Siglec F^low^ AM isolated on day 60 from IAV-infected aged mice reinfected with Adv-Control (n=4) or Adv-Tcf7l2 (n=4). K. Representative confocal microscopy IF images of KRT8 staining in lung samples isolated on day 60 from IAV-infected aged mice reinfected with Adv-Control (n=5) or Adv-Tcf7l2 (n=5). The right bar graph showing quantification of KRT8 staining in lung samples isolated on day 60 from IAV-infected aged mice reinfected with Adv-Control (n=5) or Adv-Tcf7l2 (n=5). L. Representative confocal microscopy IF images of AT1 (PDPN+) and AT2 (proSP-C+) in lung samples isolated on day 60 from IAV-infected aged mice reinfected with Adv-Control (n=5) or Adv-Tcf7l2 (n=5). Bar graph showing quantification of AT1 (PDPN+) and AT2 (proSP-C+) cells, in lung samples isolated on day 60 from IAV-infected aged mice reinfected with Adv-Control (n=5) or Adv-Tcf7l2 (n=5). M. Representative Masson trichome staining showing fibrosis in the lung section isolated on day 60 from IAV-infected aged mice reinfected with Adv-Control or Adv-Tcf7l2. N. Evaluation of lung function on day 60 from IAV-infected aged mice reinfected with Adv-Control (n=5) or Adv-Tcf7l2 (n=5). Parameters such as Respiratory system resistance (Rrs), Compliance of the Respiratory System (Crs), Newtonian Resistance (Rn), tissue resistance (G), tissue elastance (H) and respiratory system elastance (Ers), were tested on day 60. Data are representative of at least two independent experiments with similar results, *p*-values are represented as ns = non-significant, \**p*< 0.05, \*\**p*< 0.01, \*\*\**p*< 0.005, \*\*\*\**p*< 0.001.

As aged AM has higher Siglec F^low^ population, possibly derived from monocytes, we were curious to know if TCF4 over-expression can alter AM phenotype. To this end, we infected aged naive mice with Adv-Control and Adv-Tcf7l2 (Fig. 8B). Intriguingly, TCF4 over-expression diminished the Siglec F^low^ population, while increased the Siglec F^high^ population and expression in AMs of aged mice (Fig. 8C). Furthermore, frequency of Ki67^+^ AM in TCF4 over-expression was significantly improved (Fig. 8D). To further validate these findings, GFP⁺ AMs were sorted from Adv-Control or Adv-Tcf7l2 infected aged mice and analyzed for Ki67 staining and CFU potential (Fig. S8E). GFP⁺ AMs transduced with Adv-Tcf7l2 displayed higher Ki67 positivity with TCF4 co-expression and showed an enhanced capacity to form CFUs upon TCF4 overexpression (Fig. 8E, S8E). These data suggested that TCF4 over-expression promotes aged AMs to a mature embryonic-derived AM phenotype (likely due to convert those age-associated MoAM maturation)(*43*) and dampens age-associated decline in AM self-renewal.

Aging significantly increases the risk of morbidity and mortality during respiratory virus infections, including influenza and SARS-CoV-2. Taking the protective ability of TCF4 over-expression in young mice, we wondered whether similar accounted with aged mice. To this end, we infect the adenovirus transduced aged mice with IAV or SARS-CoV-2 and monitor for morbidity and mortality (Fig. 8B). TCF4 overexpression significantly improved both morbidity and mortality outcomes in aged mice following influenza infection, and reduced morbidity in the case of SARS-CoV-2 infection. (Fig. 8F-G, S8F).

Given the observed downregulation of TCF4 during chronic respiratory viral infections in both humans and mouse models, we investigated whether restoring TCF4 expression in aged mice could mitigate chronic sequelae after acute viral infection and promoted lung function. We previously reported that IAV infection in aged mice resulted in persistent lung fibrosis and chronic impairment of lung function for at least two months, which mimics chronic lung fibrosis and lung function decline observed in PASC-PF patients (*19, 44*). To this end, we infected aged mice with IAV. Then, these mice were infected with a control virus or Adv-Tcf7l2 at 28 d.p.i., two weeks after the clearance of virus infection (Fig. 8H) (*44*). We first found that TCF4 overexpression resulted in increased presence of mature AMs by 60 d.p.i (Fig. 8I-J). Furthermore, TCF4 overexpression led to a decreased accumulation of KRT8⁺ cells, indicating diminished dysplastic lung remodeling (Fig. 8K). In addition, enhanced alveolar regeneration was observed in Adv-Tcf7l2 group as evidenced by the increased numbers of AT1 and AT2 cells (Fig 8L). This was accompanied by reduced collagen deposition and lung inflammation (Fig. 8M, S8G), highlighting the potential role of the promotion of mature AMs in resolving chronic pulmonary inflammation.

To determine whether these histological improvements can be translated into better lung function, we measured lung function through FlexiVent. As shown before, IAV infection resulted in sustained defects in pulmonary function in aged mice at 60 d.p.i. following IAV infection (Fig. 8N). Respiratory System Resistance (Rrs), which reflects total airway resistance including both central and peripheral compartments (*45*), were elevated in infected aged mice, indicating airflow obstruction due to inflammation or constriction. TCF4 overexpression reduced Rrs to near-basal levels (Fig 8N). Compliance of the Respiratory System (Crs), representing the lung’s ability to expand (*46*), was markedly reduced in infected aged mice, indicative of stiff or fibrotic lungs, but was substantially restored with TCF4 overexpression (Fig 8N). Newtonian Resistance (Rn), measuring resistance in the central airways (trachea and bronchi) (*47*), was also elevated in infected aged mice and significantly reduced by TCF4 over-expression (Fig 8N), suggesting relief of bronchoconstriction. We further assessed Tissue Resistance (G), and Tissue Elastance (H), reflecting lung tissue stiffness or recoil (*48*), which was elevated in aged-infected lungs but significantly lowered in TCF4-treated mice(Fig 8N), suggesting reduced fibrotic tissue accumulation. Finally, Respiratory System Elastance (Ers), the inverse of Crs, measuring overall lung and chest wall stiffness was improved in TCF4-overexpressing mice, indicating enhanced lung flexibility (Fig 8N). Together, these data demonstrate that TCF4 overexpression significantly improves key lung function parameters, highlighting the therapeutic potential in targeting a key macrophage stem factor to diminish chronic lung fibrosis and improve lung function post-acute viral infection including PASC-PF.

## Discussion

Our study identifies the conventional Wnt-family transcription factor TCF4 (Tcf7l2) as a master regulator of AM stemness, metabolic fitness and reparative function. By combining genetic loss- and gain-of-function approaches with acute and chronic influenza and SARS-CoV-2 infection models, we demonstrate that TCF4 is indispensable for the developmental maturation of AMs, their self-renewal capacity and AM-mediated effective host recovery after severe viral pneumonia. Conversely, enforced expression of TCF4 promotes AM proliferation, limits acute lung injury, and attenuates post-viral fibrosis, even in aged hosts. These data reveal an actionable pathway for treating acute and chronic lung diseases, including the debilitating pulmonary conditions associated with long COVID.

In contrast to the classical view of TCF/LEF proteins as nuclear co-effectors of β-catenin (*49*) our data reveal a strikingly different arrangement in AMs, in which TCF4 and β-catenin function as antagonistic partners. In the absence of TCF4, AMs downregulate classical maturation factors, upregulate repressors of self-renewal like *Mafb* and *c-Maf* (*2*), and transition to a pro-inflammatory state driven by β-catenin-HIF-1α activation (*11*). Inhibition of β-catenin, in turn, upregulates TCF4, whereas TCF4 overexpression suppresses Wnt target genes and β-catenin signaling. Interestingly, TCF4 and β-catenin can occupy the same regulatory regions of key metabolic genes, indicating a potential model of direct competition for promoter access. This competitive binding likely dictates whether the cell prioritizes mitochondrial metabolism and self-renewal (driven by TCF4) or glycolysis-fueled inflammation (driven by β-catenin/HIF-1α). This unique regulatory logic may be a specialized adaptation in long-lived, self-renewing AMs to maintain immune quiescence and tissue integrity in the delicate alveolar oxygen-rich niche. Whether this TCF4/β-catenin antagonistic relationship is unique to AMs or represents a broader paradigm in other tissue-resident macrophages such as microglia and Kupfer cells warrants further investigations.

Our findings also provide a molecular explanation for the vulnerability and dysfunction of AMs during severe respiratory viral infections (*50, 51*). We show that both influenza and SARS-CoV-2 infections potently suppress TCF4 expression at the peak of inflammation, a process mediated largely by exuberant interferon signaling, particularly IFN-γ. While TCF4 expression is restored following uncomplicated infections, its suppression persists in hosts with chronic tissue sequelae including PASC. We speculate that sustained TCF4 loss creates a feed-forward loop of AM impairment, MoAM recruitment, resulting accumulation of dysplastic KRT8^hi^ transitional epithelial cells, a hallmark of post-viral fibrosis in mice and humans (*18, 19, 21*). Thus, failure to re-establish the TCF4-dependent mature AM gene program provides a mechanistic explanation for the defective alveolar repair observed in PASC-PF lungs (*41*).

Aging is associated with increased susceptibility to respiratory viral infections and impaired recovery, contributing to a higher risk of developing PASC (*52*). Aging also profoundly impairs AM self-renewal and promotes the establishment of immature MoAM (*42, 53, 54*). We found that AMs from aged mice exhibit a baseline deficiency in TCF4 and Ki-67 expression, mirroring the erosion of stemness seen during acute infection. Remarkably, adenoviral delivery of TCF4 to aged mice restored a mature, embryonic-derived macrophage-like, Siglec F^high^ AM compartment and re-engaged their proliferative capacity. When these aged animals were challenged with influenza or SARS-CoV-2, the enforced TCF4 expression blunted weight loss, improved survival, and preserved alveolar architecture. Importantly, TCF4 overexpression-mediated benefits on host recovery were contingent on an intact AM pool as clodronate-mediated macrophage depletion abolished the protective effect. Thus, our data strongly indicates that the restoration of the AM compartment, particularly their replicative ability, is a viable strategy to reverse age-associated vulnerability to viral disease.

Most strikingly, post-acute administration of TCF4, initiated weeks after viral clearance, reversed established hallmarks of chronic lung disease in aged mice. This intervention curtailed the persistence of aberrant KRT8^hi^ epithelial cells, reduced collagen deposition, and promoted alveolar regeneration. Importantly, as confirmed by FlexiVent analysis, these structural improvements translated directly to the restoration of pulmonary function. This demonstrates that TCF4 supplementation not only prevents but can also repair age-associated defects in alveolar regeneration. Given that treatments for long COVID and other post-viral syndromes remain largely supportive currently (*16, 55, 56*), our findings reveal the TCF4 axis as an actionable target that can simultaneously dampen inflammation and boost tissue regeneration. These proof-of-concept data suggest that next-generation therapeutics, such as small-molecule agonists or cell-based strategies designed to enhance TCF4 activity, could potentially complement antiviral and anti-fibrotic regimens to dampen chronic diseases after acute infection. These findings broaden the therapeutic potential of TCF4 modulation beyond acute viral pneumonia to encompass the treatment of age-related chronic pulmonary fibrosis, a condition for which effective therapies remain limited currently.

### Limitations of the study

First, the precise molecular signals by which IFN signaling represses TCF4 transcription warrants further investigation. Identifying the specific transcription and epigenetic factors involved could reveal additional therapeutic targets. Second, currently there is a lack of AM-specific, particularly inducible Cre line, that enables to acutely and selectively deplete TCF4 in AMs. Due to this limitation, we had to use the Lyz2-cre ERT2 strategy to inducible ablate TCF4 in myeloid cells. Therefore, we could not completely exclude the effects of TCF4 depletion and overexpression in other myeloid cells *in vivo* on host recovery. Nevertheless, the low expression of TCF4 in monocytes and neutrophils coupled with mixed bone marrow chimera and *in vitro* experiments strongly suggest that the intrinsic role of TCF4 on AM phenotypes. Finally, while our adenoviral approach demonstrates proof-of-concept, developing clinically translatable strategies to modulate TCF4 activity in the lung, such as small molecule agonists or targeted gene therapy, will be a critical next step.

## Methods

### Ethics and Biosafety

This study was approved by the Institutional Review Board (IRB) at Cedars-Sinai Medical Center (IRB# Pro00035409) and the University of Virginia (IRB# 13166). All animal experiments were conducted in accordance with protocols approved by the Institutional Animal Care and Use Committees (IACUC) of the University of Virginia (Charlottesville, VA) otherwise specified, adult mice used in experiments were age- and sex-matched. All SARS-CoV-2-related work was performed in the ABSL-3 facilities at the University of Virginia, while experiments involving influenza virus were carried out in ABSL-2 facilities at University of Virginia.

### Mice

The following mouse strains were obtained from The Jackson Laboratory: WT C57BL/6 (Cat# 000664), *Lyz2*-Cre (Cat# 004781) and *Lyz2-Cre* ERT2 (Strain #:031674). All strains were subsequently bred in-house under specific pathogen-free (SPF) conditions. Tcf7l2^cKO^ and Tcf7l2^icKO^ were generated by crossing Tcf7l2^fl/fl^ mice with Lyz2-Cre or Lyz2-ERT2-Cre mice, respectively.

### Viruses

The Influenza H1N1 A/PR8/34 virus was propagated in the allantoic cavity of 10-day-old embryonated hen’s eggs and confirmed to be free of bacterial, mycoplasma, and endotoxin contamination. The isolated virus stocks were stored at −80 °C and titrated using Madin–Darby canine kidney (MDCK) cells. The mouse-adapted SARS-CoV-2 strains, MA30 kindly provided by Dr. Stanley Perlman (University of Iowa) and Dr. Barbara J. Mann (University of Virginia School of Medicine), respectively. These strains were cultured in Vero E6 cells (ATCC CRL-1587), and viral titters were determined using a plaque assay on Vero E6 cells.

### Infection Protocol

Wild-type (WT) C57BL/6 mice (8–12 weeks old), Tcf7l2^fl/fl^, Tcf7l2^cKO^, Tcf7l2^icKO^ mice were infected with 150 PFU influenza A virus (IAV) otherwise mentioned. Middle-aged and advanced aged were infected with 75 PFU IAV for morbidity assessments or 150 PFU for mortality studies. For SARS-CoV-2 (MA30 strain) infection, Tcf7l2^icKO^ (8–12 weeks old) were inoculated with 2000 PFU to assess morbidity. For adenovirus transduction 5X10^4^ PFU were used. All infections were administered intranasally under anesthesia. Mice were weighed before infection to establish a baseline, followed by daily weight monitoring. A 30% weight loss relative to baseline was designated as the clinical endpoint.

### Mouse Lung Tissue Processing and BAL Cell Collection

BAL was collected by flushing the airway three times with 600 µL of sterile PBS via a tracheal incision. BAL cells were pelleted by centrifugation for flow cytometry analysis, AM sorting, and immunofluorescence (IF) staining. The BAL supernatants were stored for cytokine/chemokine quantification, protein concentration analysis. Following BAL collection, the right ventricle of the heart was perfused with 10 mL of chilled 1X PBS. The right lung lobes were excised, finely minced, and digested using 180 U/mL Collagenase Type 2 (Worthington Biochemical) for 30 minutes at 37°C. Tissue was then mechanically disrupted using a gentleMACS tissue dissociator (Miltenyi). Single-cell suspensions were prepared by red blood cell lysis, filtration through a 70 μm mesh, and washing with flow cytometry buffer. The resulting cell suspension was centrifuged and resuspended in an appropriate buffer for flow cytometry analysis.

For histological analysis, 10% paraformaldehyde (PF) was gently instilled into the left lung lobe, keeping it inflated for one minute before excision. The lobe was then fixed in 10% PF for 48 hours, followed by transfer to 70% ethanol. Samples were sent to the UVA Research Histology Core Facility for paraffin embedding, and 5 µm sections were prepared for Hematoxylin and Eosin (H&E) staining, Masson’s trichrome staining, and immunofluorescence analysis.

Slides were digitally scanned by the UVA Biorepository and Tissue Research Facility Core and further processed using ImageJ (version 1.54). Areas of inflammation were manually annotated, and a random trees classifier was interactively trained to distinguish inflammatory regions from surrounding tissue. This classifier employed machine learning techniques to accurately segment and classify various tissue components.

### Mouse Alveolar Macrophage (AM) Culture and *In vitro* Treatment

Mouse AMs were isolated from BAL as previously described. For naïve mice, AMs were purified by adherence for 2 hours in complete culture medium (RPMI 1640 supplemented with 10% FBS and 1% penicillin/streptomycin/glutamate) at 37°C and 5% CO₂. Nonadherent cells were removed by washing with warm PBS, and adherent AMs were maintained in complete medium supplemented with 10 ng/mL recombinant murine GM-CSF (BioLegend, catalog no. 576304). For *in vitro* stimulation, AMs were treated with the following reagents at the indicated concentrations for 24h unless otherwise noted: R848 (50 ng/mL; Invivogen, tlrl-r848), CpG (5 μg/mL; Invivogen, tlrl-1826), LPS (20 ng/mL; Invivogen, tlrl-eklps), IL-6 (20 ng/mL; BioLegend, 575704), IL-1β (20 ng/mL; BioLegend, 575104), TNFα (25 ng/mL; PeproTech, 315-01A), IFNγ (50 ng/mL; PeproTech, 315-05), IFNα (50 ng/mL; BioLegend, 752804), IFNλ (50 ng/mL; BioLegend, 575304), or Poly(I:C) (5 μg/mL; Invivogen, tlrl-pic).

### Bulk RNA Sequencing

Total RNA was extracted from *in vitro* cultured alveolar macrophages (AMs) for bulk RNA sequencing. Following quality assessment, only high-quality RNA samples (RIN > 7.0; Agilent Bioanalyzer) were used for library preparation. cDNA synthesis, end repair, A-tailing, and ligation of Illumina-indexed adapters were carried out using the TruSeq RNA Sample Prep Kit v2 (Illumina, San Diego, CA), according to the manufacturer’s protocol. Library quality and concentration were assessed using the Agilent Bioanalyzer DNA 1000 chip (Santa Clara, CA) and Qubit fluorometry (Invitrogen, Carlsbad, CA). Paired-end sequencing was performed on the Illumina HiSeq 4000 platform using the Illumina cBot and HiSeq 3000/4000 PE Cluster Kit, following standard Illumina protocols. Base calling was conducted with Illumina’s RTA software (v2.5.2). Reads were aligned to the mouse reference genome (GRCm38/mm10) using the spliced-read aligner Tophat2 (v2.2.1). Pre- and post-alignment quality control, gene-level quantification, and normalization (FPKM) were performed using the RSeQC package (v2.3.6), with annotation from the NCBI mouse RefSeq gene model. Differential gene expression between experimental groups was determined using DESeq2 (Wald test). For data visualization, expression values were log-transformed, and genes showing a log2 fold change > 2 and log10(P-value) > 25 were highlighted. Gene Set Enrichment Analysis (GSEA) was performed using gene sets from the MSigDB database, employing a weighted enrichment statistic and log2 ratio ranking metric.

### Bone Marrow Chimeras

Bone marrow cells from CD45.1 WT and Tcf7l2^cKO^ were mixed at a ratio of 1:1, the mixed bone marrow cells were then transferred to the irradiated (1,100 rad) WT CD45.2+ recipient mice to create chimeric mice. After 8 weeks’ reconstitution, chimeric mice were used for isolation of BAL fluid to study AM populations. Additionally, blood was collected to ensure equal transfer on bone marrow cells.

### Human Lung Tissue Specimens

Human lung samples were collected from patients enrolled in the IRB-approved Lung Institute BioBank (LIBB) studies at Cedars-Sinai Medical Center, Los Angeles, CA, and the University of Virginia, Charlottesville, VA. Informed written consent was obtained from all participants or their legal representatives. Lung tissues were processed within 24 hours of surgical excision. Tissues were sectioned and immediately fixed in 10% neutral-buffered formalin for 24 hours, followed by automated processing using the HistoCore PEARL Tissue Processor (Leica) and paraffin embedding for histological analysis. The formalin-fixed, paraffin-embedded (FFPE) blocks were stored at room temperature until further sectioning and use.

### Cell Immunofluorescence

For *in vitro* experiments, AMs were cultured and treated on coverslips. Cells were fixed using 4% paraformaldehyde in PBS for 10 min. Afterward, they were washed in PBS and permeabilized with 0.1% Triton X-100 in PBS for 15 min. Subsequently, cells were incubated with the primary antibody in Agilent Dako antibody dilute solution (S302283) overnight at 4°C. Following PBS washes, secondary antibodies (1:500) were applied and incubated for 1 hour at room temperature. Coverslips were then mounted onto glass microscope slides using ProLong Gold Antifade Mountant with DAPI (Invitrogen). The following antibodies were used TCF4 (Santa Cruz Biotech sc-166699, 1:200), Ki67 (Thermo, Ki-67, (eBioscience™ Cat #14-5698-82), Non-phospho (Active) β-Catenin (Cell Signaling Tech Cat#8814 and HIF-1α (Cell Signaling Tech Cat #79233).

### Lung Immunofluorescence

Paraffin-embedded lung tissue sections were deparaffinized in xylene and rehydrated through a graded ethanol series. Antigen retrieval was performed using either 1× Agilent Dako Target Retrieval Solution (pH 9.0; Cat# S236784) or sodium citrate buffer (10 mM sodium citrate, 0.05% Tween-20, pH 6.0). Retrieval was conducted in a steamer for 20 minutes for mouse lungs and 45 minutes for human lungs. Following antigen retrieval, sections were blocked and surface stained. For intracellular targets, tissues were permeabilized using 0.5% Triton X-100 with 0.05% Tween-20 for 1 hour at room temperature.

Sections were incubated with primary antibodies overnight at 4 °C. After washing, fluorescently labeled secondary antibodies were applied for 2 hours at room temperature. Nuclei were counterstained with DAPI (1:5000; ThermoFisher Scientific) for 3 minutes. Slides were mounted using ProLong Diamond Antifade Mountant (ThermoFisher Scientific) and cured at room temperature for 24 hours. Imaging was performed using Zeiss confocal microscope. Image processing and analysis were conducted using ImageJ Fiji, and Zen 3.5 Blue software (Zeiss). Following antibodies were used, TCF4 (Santa Cruz Biotech, Mouse sc-166699, 1:200), CD68 Rabbit mAb #76437, 1:200), SiglecF Rat (Thermo Cat # 14-1702-82, 1:200), cytokeratin 8 Rat (TROMA-1, DSHB, AB_531826, 1:300), MERTK Rat (eBioscience Cat # 14-5751-82 ). PDPN

Hamster (Abcam Cat#: ab11936, 1:300), proSP-C Mouse (Santa Cruz Cat#: sc-518029, 1:300). Secondary antibody such as Goat anti-mouse IgG AF488 (ThermoFisher, A-11001, 1:300), Goat anti-rat IgG AF555 (ThermoFisher, A48270, 1:300), Goat anti-Armenian Hamster IgG AF488 (ThermoFisher, A78963,1:400), Goat anti-rat IgG AF647(ThermoFisher, A-21247, 1:200), Goat anti-rabbit IgG AF488 (ThermoFisher, A-11008, 1:300), were used.

### RNA Isolation and Quantitative PCR with Reverse Transcription

Total RNA from AMs and bone marrow-derived AM were extracted using the RNeasy Kit (Qiagen) following the manufacturer’s instructions. Reverse transcription was performed using M-MLV Reverse Transcriptase (200 U/μL) (Invitrogen) as per the manufacturer’s guidelines. Quantitative PCR (qPCR) was conducted using PowerUp Sybr Green (Life Technologies). Cq values were obtained using a CFX96 PCR system (Bio-Rad) and analyzed using the ΔCq method, normalizing target gene expression to *Gapdh*. Primer sequences are provided in Supplementary Table 1.

### Immunoprecipitation and Immunoblots

Cells were lysed in Cell Lysis Buffer (Cell Signaling Tech, Cat# #9803) supplemented with a protease inhibitor cocktail (Sigma, Cat# P8340) and protein concentrations were quantified using the BCA protein assay kit. Proteins were separated on a 4–12% SDS-PAGE gel. GAPDH (Cell Signaling Technology, Cat# 97166S, 1:2000), β-Actin (Cell Signaling Technology Cat #4967 1:2000), TCF4 (Santa Cruz Biotech sc-166699, 1:2000), β-Catenin (Cell Signaling Tech Cat #8480 1:2000), and HIF-1α (Cell Signaling Tech Cat #79233 1:2000).

### Flow cytometry

Single cell suspensions were preincubated with anti-FcgRIII/II (Fc block) before 30 min of incubation with appropriate fluorochrome-labelled antibodies. The following antibodies (supplied by BioLegend or BD Biosciences) and staining reagents were used: CD45-PerCP/Cy5.5 (BioLegend, clone 30-F11, catalog no. 103132), Siglec-F-BV421 (BD Biosciences, clone E50-2440, catalog no. 562681), CD11b–FITC (BioLegend, clone M1/70, catalog no. 101206), CD11c-BV510 (BioLegend, clone N418, catalog no. 117338), Ly6G–PE/Cy7 (BioLegend, clone 1A8, catalog no. 127618), Ly6C-BV711 (BioLegend, clone HK1.4, catalog no. 128037), CD64-PE (BioLegend, clone X54-5/7.1, catalog no. 139304), MerTK–APC (BioLegend, clone 2B10C42, catalog no. 151508), The dilution of surface staining Abs was 1:200. Staining samples were analyzed on a FACS Attune or FACS Attune NXT flow cytometer (Life Technologies) and interpreted using the software FlowJo (Treestar).

### BCA Protein Assay

Protein concentration was measured using the BCA Protein Assay Kit (Thermo Scientific, Cat# 23225). For each bronchoalveolar lavage (BAL) sample, 2 μL was used in the assay. Colorimetric quantification was performed using a VERSAmax microplate reader (Molecular Devices) at a wavelength of 570 nm.

### Tamoxifen treatment

To induce gene deletion in Tcf7l2^icKO^ mice, tamoxifen (Sigma) was dissolved in 0.5 ml ethanol and 9.5 ml warm sunflower oil (Sigma-Aldrich). The suspension was administered at 2 mg/mouse at a concentration of 20 mg/ml via i.p. injection for 5 consecutive days.

### Administration of clodronate liposomes (CL)

CL were obtained from Encapsula NanoSciences. Mice were depleted of AM by intranasally administering 50 µL of clodronate liposomes (CL) for 2 consecutive days. The mice were given two-day rest prior to influenza A virus (IAV) infection. Control liposome was used as control.

### CFU assay

Freshly isolated AM were seeded at a density of 20,000 cells per 35 mm culture dish and allowed to adhere for 2 hr at 37°C with 5% CO₂ Following incubation, cells were washed with PBS and subsequently cultured in MethoCult medium (M3231, Stem Cell Technologies) supplemented with 10 ng/ml GM-CSF, 50 μg/ml penicillin/streptomycin, and 2 mM glutamine. Colony formation was assessed on day 18 post-plating.

### Metabolic Analysis

Real-time measurements of oxygen consumption rate (OCR) and extracellular acidification rate (ECAR) in alveolar macrophages (AMs) were performed using a Seahorse XFp Analyzer (Seahorse Bioscience). AMs (1 × 10⁵ cells per well) were seeded into Seahorse XFp Cell Culture Miniplates and pre-treated overnight at 37°C with 5% CO₂ in the presence or absence of 100 ng/ml Wnt3a. The following day, cells were washed twice and incubated at 37°C for 1 hour in unbuffered assay medium (pH 7.4, Agilent Technologies) without CO₂. For the mitochondrial stress test, the medium contained 10 mM glucose, while for the glycolytic stress test, glucose was omitted. OCR and ECAR were recorded under basal conditions and in response to sequential additions of metabolic inhibitors: 1 μM oligomycin, 1.5 μM FCCP (carbonyl cyanide-4-(trifluoromethoxy) phenylhydrazone), 0.5 μM rotenone + 0.5 μM antimycin, 10 mM glucose, and 50 mM 2-DG (2-deoxy-D-glucose) (all from Sigma). Data was analyzed using Wave Desktop software version 2.6 (Agilent Technologies).

### Single cell RNA sequencing

For library preparation, lungs were isolated from Tcf7l2^fl/fl^ or Tcf7l2^icKO^ mice on day 9 following tamoxifen treatment from day 4. Within each group single cells were isolated from lungs and were pooled and labeled with BioLegend TotalSeq™-A antibodies according to the manufacturer’s protocol. Pooled hash-tagged libraries were generated using the 10x Genomics Chromium Next GEM Single Cell 3′ Reagent Kit v3.1 (Dual Index). Sequencing was performed on the NovaSeq X Plus platform (Admera Health). Raw sequencing data were processed and aligned using Cell Ranger. The curated data were subsequently subjected to demultiplexing, quality control, filtering, differential gene expression analysis, and cluster identification using the Seurat (v5.1.0) package in R (v4.3.3). Downstream gene set enrichment analysis was conducted with the clusterProfiler (v4.10.1) package, using gene sets obtained from MSigDB.

### Assessment of Respiratory Mechanics and Lung Function

Lung function was assessed using the forced oscillation technique as previously described. Briefly, mice were deeply anesthetized with an intraperitoneal injection of ketamine (100 mg/kg) and xylazine (10 mg/kg), followed by administration of pancuronium bromide (1 mg/kg, intraperitoneally) to induce paralysis and eliminate spontaneous respiratory efforts. A tracheostomy was performed using a blunt 18-gauge cannula (resistance: 0.18 cm H₂O·s·ml⁻¹), which was secured with a nylon suture. Mice were then connected to a computer-controlled ventilator (flexiVent, SCIREQ), and respiratory mechanics were measured under tidal breathing with a positive end-expiratory pressure (PEEP) of 3 cm H₂O.

To ensure accurate assessment, closed airways were recruited prior to measurements by delivering two rapid inflations to total lung capacity, each maintained for several seconds to simulate a deep breath. Baseline measurements were normalized to each animal’s maximal lung capacity, allowing for the detection of both large and small airway dysfunction during normal (tidal) breathing. Only data sets with >90% model fit accuracy, as determined by the software’s constant-phase model, were included in the analysis. Throughout the procedure, a heart rate of approximately 60 beats per minute confirmed that the animals remained alive during data collection.

### Statistical analysis

Data means SEM of values from individual mice (*in vivo* experiments). Unpaired two-tailed Student’s t test (two group comparison), one-way ANOVA (multiple group comparison), Multiple t tests were used to determine statistical significance by GraphPad Prism software. We consider P values < 0.05 as significant. ∗, p < 0.05; ∗∗, p < 0.01; ∗∗∗, p < 0.001.

**Table 1.**
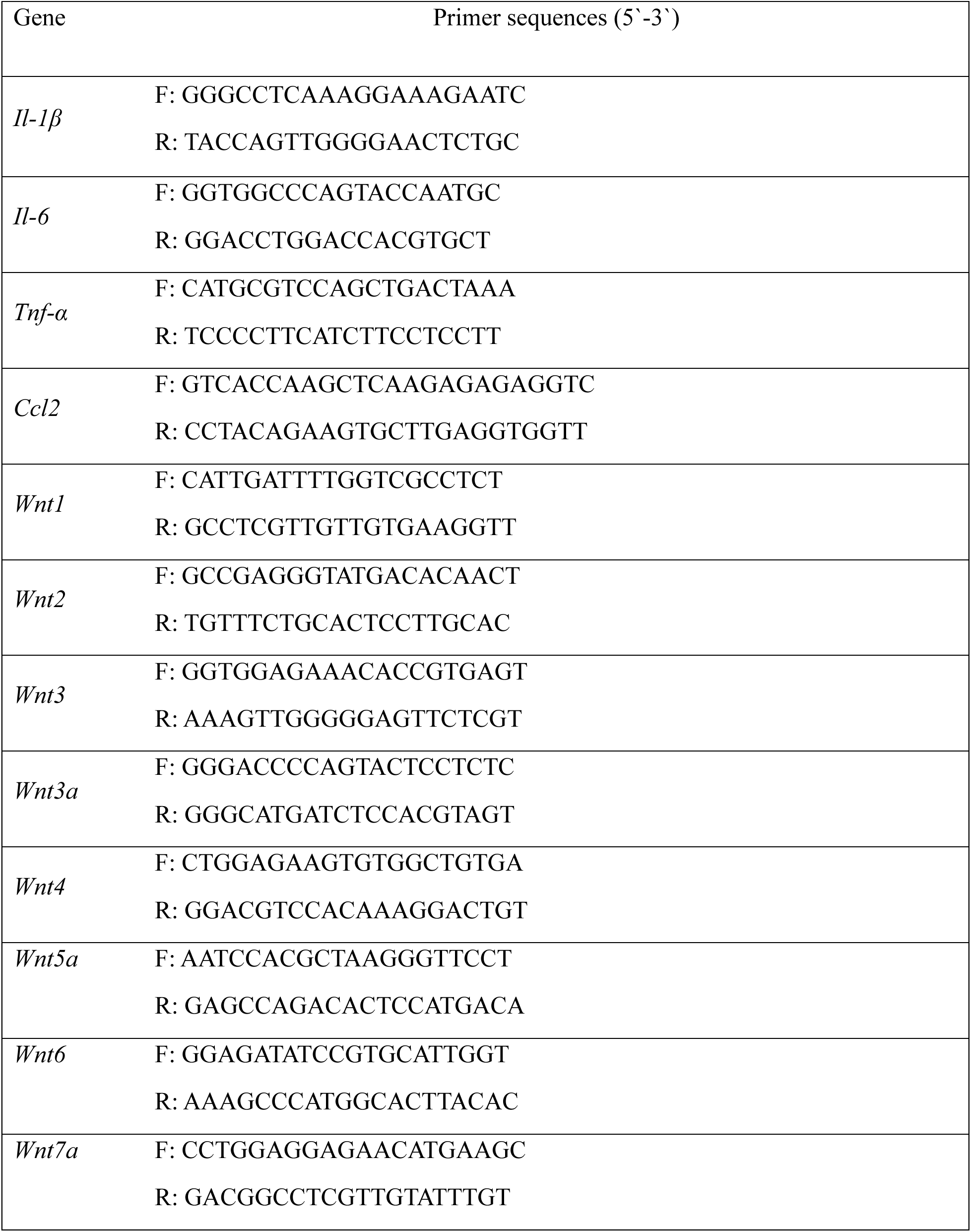
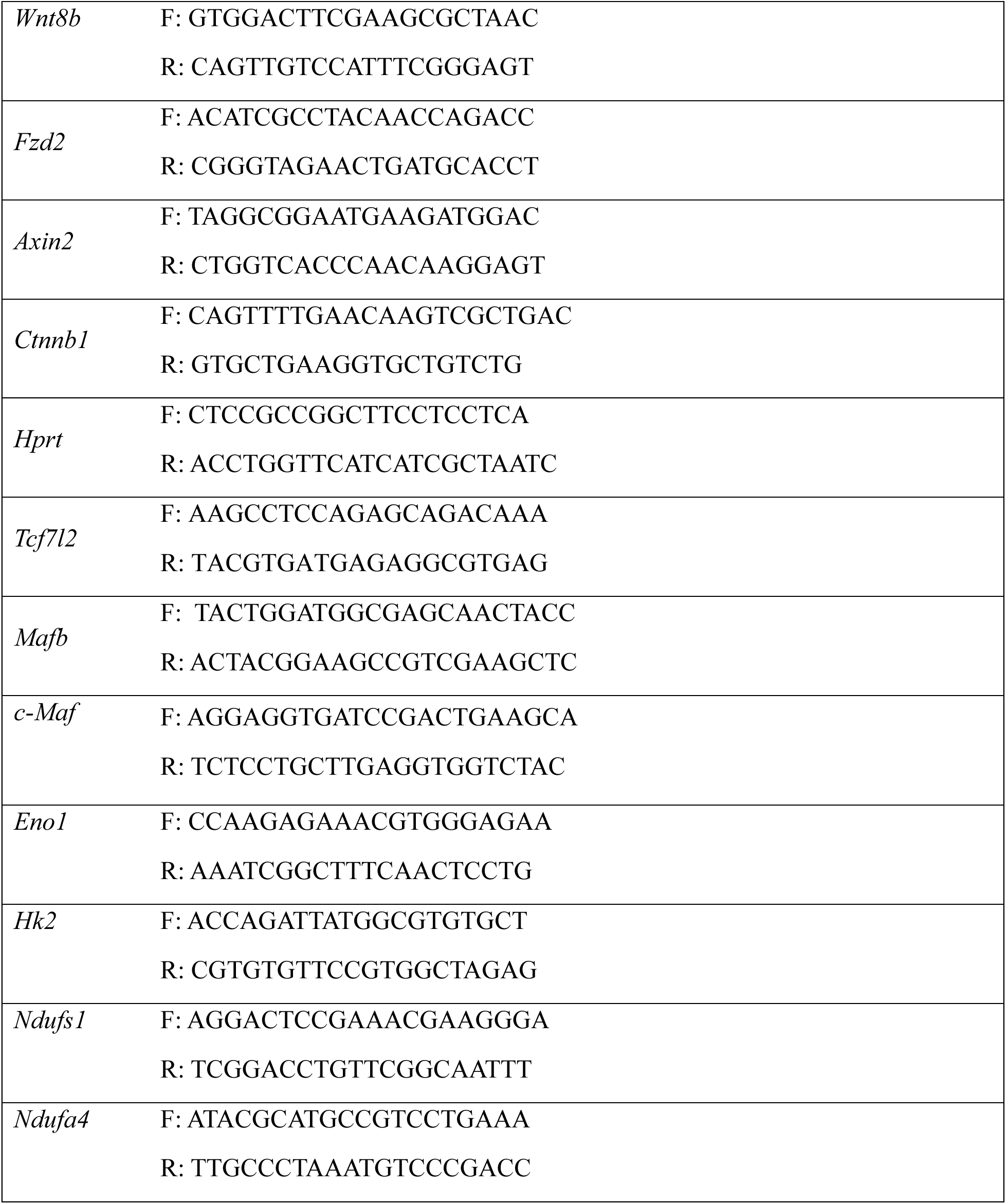
qRT-PCR Primers.

## Acknowledgement

We gratefully acknowledge the Lung Institute BioBank staff at Cedars-Sinai Medical Center for their support in providing human lung specimens. We appreciate the assistance of Drs. Jay Kolls and Nick Maness with virus propagation. The schematics presented in the manuscript were created using BioRender.com. Data for this study were generated using the Flow Cytometry Core, Research Histology Core Facilities, Biorepository and Tissue Research Facility, and the Biomolecular Analysis Facility at the University of Virginia. This work was supported in part by grants from the US National Institutes of Health: AI147394, AG069264, AI112844, HL170961, AI176171, and AI154598 (to J.S.); R01HL132287, R01HL167202, and R01HL132177 (to Y.M.S.); F31HL170746 and T32AI007496 (to H.N.); R01HL155759 and R01HL159953 (to P.C.); P01-HL108793 (to D.J.); and HL159675, HL152293, and AI163753 (to J.Q.).

## Author contribution

M.A., A.S.C & J.S., conceived the overall project. M.A., A.S.C & J.S., designed the experimental strategy and analyzed data. J.T., H.N., A.F., J.T., X.Q., C.L., I.S.C., W.Q., F.N, G.A.S., S.Y.P., T.P., Y.M.S., R. V., P.C., J.S., performed experiments, analyzed data, or contributed critical reagents to the study. M.A & J.S. wrote the original draft. All authors read, edited, and approved the final manuscript.

## Supplementary Figures

**Fig. S1.**
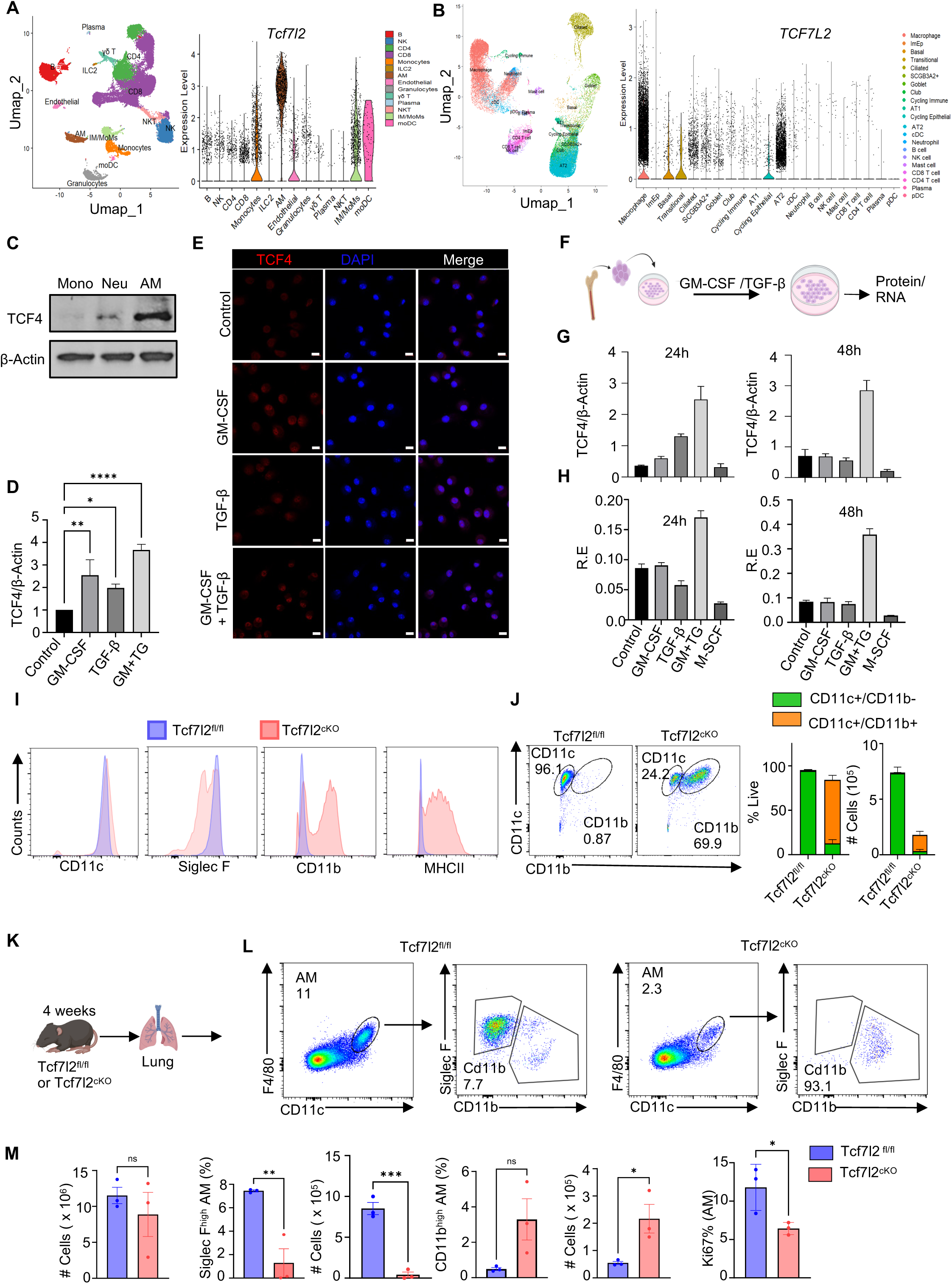
TCF4 is required for AM maturation during development. A. UMAP analysis of *Tcf7l2* expression in lung immune cells in WT mice. Violin plot of *Tcf7l2* expression in lung immune cells in WT mice. B. UMAP analysis of *TCF7L2* expression in lung immune cells in humans. Data derived from publicly available dataset (GSE224955). Violin plot of *TCF7L2* expression in lung immune cells in humans. C. Western blot showing TCF4 expression in myeloid cells such as monocytes (Mono), Neutrophils (Neu), and AM. D. Quantification of TCF4 expression in AM stimulated with or without for GM-CSF and/or TGF-β for 48h *in vitro*. E. Confocal imaging of TCF4 and DAPI in AM stimulated with or without for GM-CSF and/or TGF-β for 48h. F. Schematic diagram of monocyte derived from bone-marrow stimulated with or without for GM-CSF and/or TGF-β for 24h or 48h *in vitro*. G. Quantification of Western blot for TCF4 expression in bone-marrow derived monocytes stimulated with or without for GM-CSF and/or TGF-β for 24h or 48h *in vitro*. H. Quantification relative gene expression (R.E) for *Tcf7l2* in bone-marrow derived monocytes stimulated with or without for GM-CSF and/or TGF-β for 24h or 48h *in vitro*. I. Histogram plot for the expression of CD11c, Siglec F, CD11b, and MHCII on AM isolated from naïve Tcf7l2^fl/fl^ or Tcf7l2^cKO^ mice. J. Flow cytometry plot showing expression of CD11c^+^ and CD11b^+^ AMs isolated from Tcf7l2^fl/fl^ or Tcf7l2^cKO^ mice. Bar graph showing frequency and cell number of CD11c^+^ and CD11b^+^ AMs isolated from Tcf7l2^fl/fl^ or Tcf7l2^cKO^ mice. K. Schematic figure showing Tcf7l2^fl/fl^ and Tcf7l2^cKO^ juvenile mouse lung isolation model. L. Flow cytometry flow showing AM characterization from lung of Tcf7l2^fl/fl^ and Tcf7l2^cKO^ juvenile mice M. Bar graph showing total cell number in BAL fluid, frequency and count of Siglec F^high^, frequency and count of CD11b^+^ AM, and frequency of Ki67^+^ AMs in naïve Tcf7l2^fl/fl^ or Tcf7l2^cKO^ juvenile (4 weeks) mice. Data are representative of at least two independent experiments with similar results except A-C, *p*-values are represented as ns = non-significant, \**p*< 0.05, \*\**p*< 0.01, \*\*\**p*< 0.005, \*\*\*\**p*< 0.001.

**Fig. S2.**
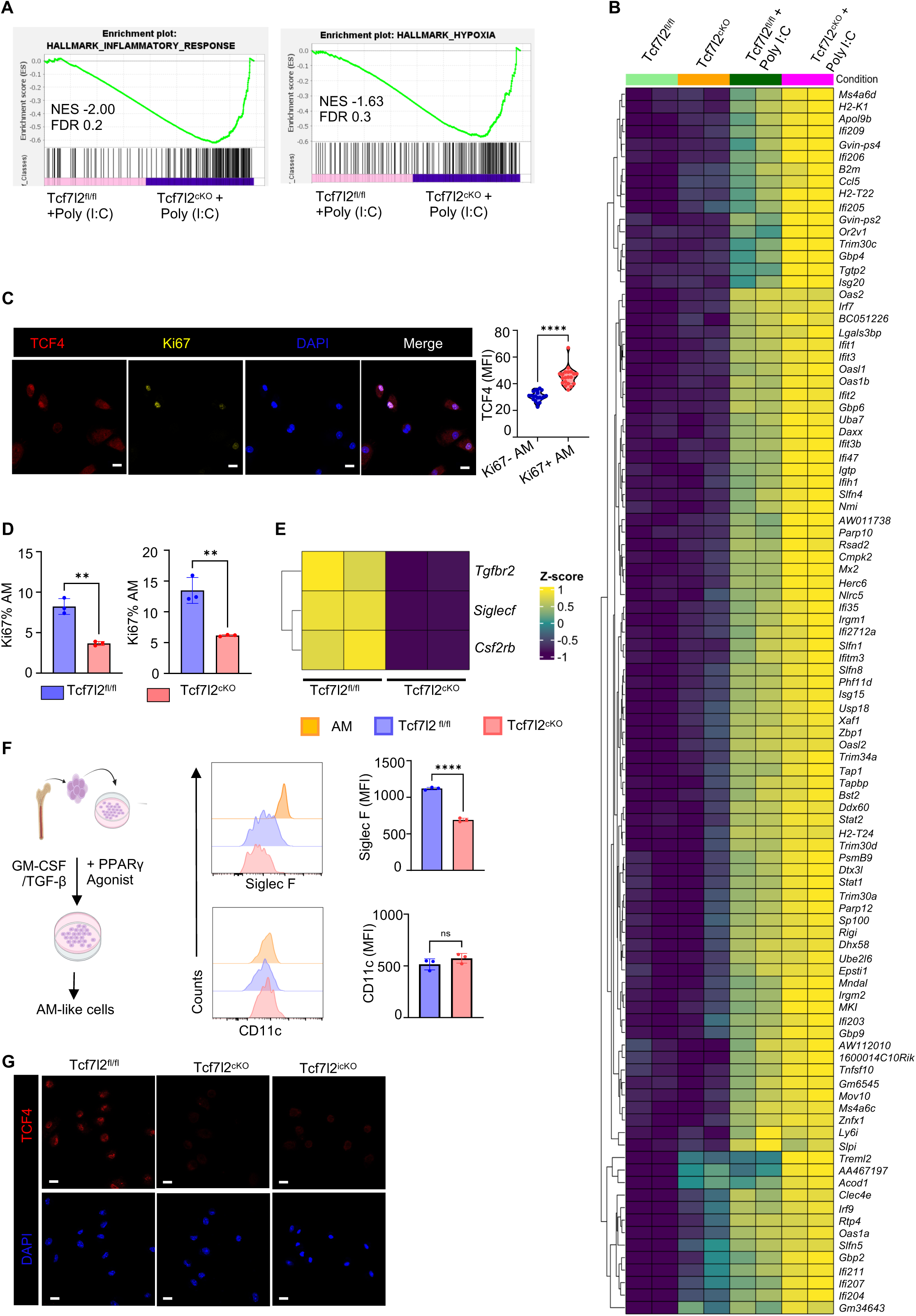
TCF4 promotes AM cell proliferation. A. Enrichment plot showing inflammatory and hypoxia related gene set in Tcf7l2^fl/fl^ or Tcf7l2^cKO^ AMs treated with or without Poly(I:C). B. Heatmap showing differential gene expression in Tcf7l2^fl/fl^ orTcf7l2^cKO^ AMs treated with or without Poly(I:C) for 24h *in vitro*. C. Representative confocal images for TCF4 and Ki67 colocalization in WT AM treated with GM-CSF and TGF-β for 48h *in vitro*. Violin plot showing TCF4 expression in Ki67^−^ and Ki67^+^ WT AMs. D. Bar graph showing Ki67^+^ in Tcf7l2^fl/fl^ or Tcf7l2^cKO^ AM stimulated with or without GM-CSF for 24h. E. Heat map showing gene expression of *Tgfbr2*, *Siglecf* and *Csf2rb* in WT and F. Representation of generation of AM-like cells from BM of WT mice. Right histogram showing Siglec F and CD11c expression in AM and AM-like cells generate from Tcf7l2^fl/fl^ or Tcf7l2^cKO^ BM cells. G. Representative confocal images showing TCF4 deletion in AM isolated from Tcf7l2^cKO^ or Tcf7l2^icKO^ as compared to Tcf7l2^fl/fl^ mice. Scale bar 10 μm. Data are representative of at least two independent experiments with similar results except A-B and E, *p-*values are represented as \*\**p*< 0.01, \*\*\*\**p*< 0.001.

**Fig. S3.**
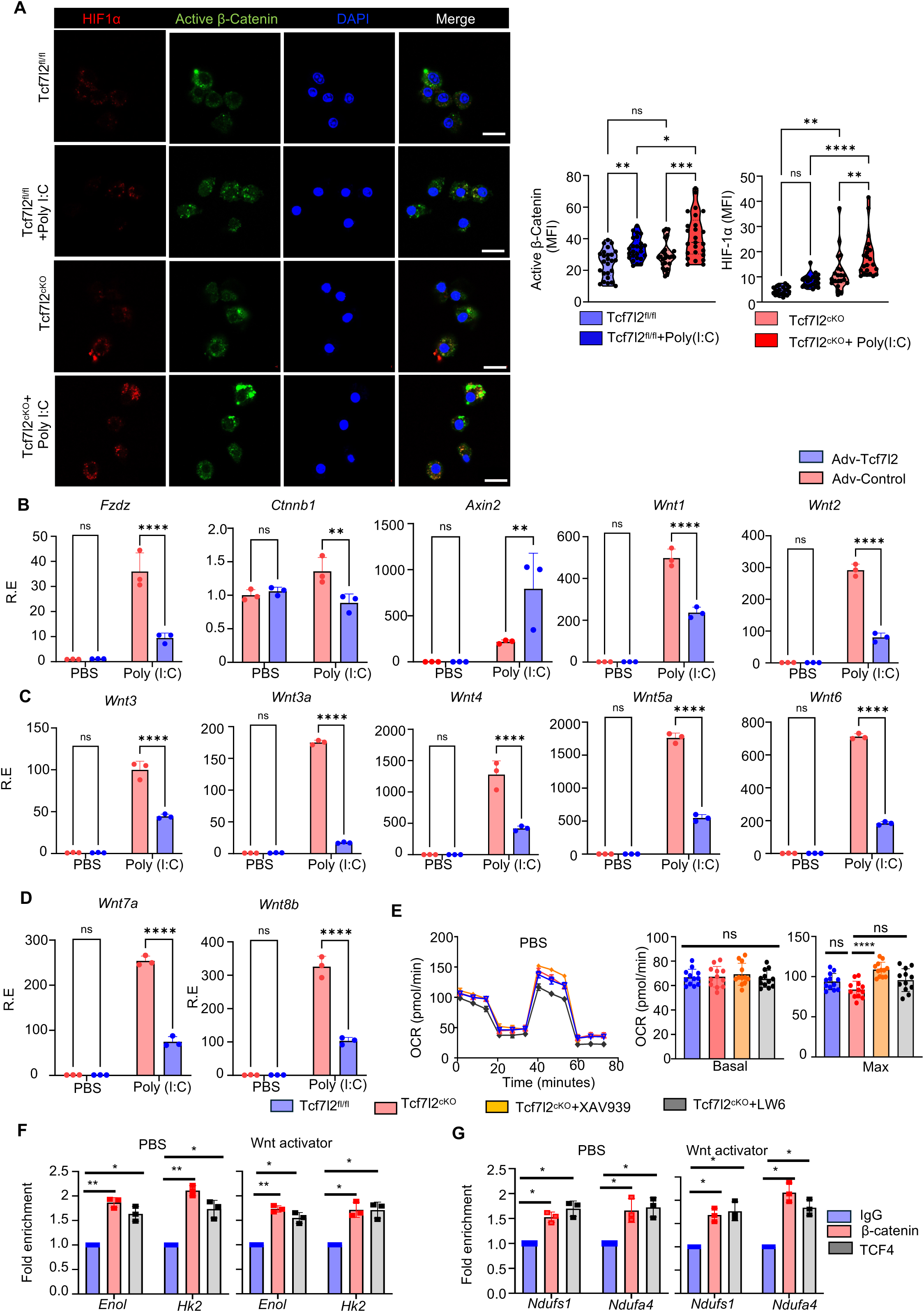
TCF4 controls Wnt-β-catenin signaling. A. Representative confocal images of Tcf7l2^fl/fl^ or Tcf7l2^cKO^ AM stained with HIF-1α and active β-catenin, stimulated with Poly(I:C) for 24h *in vitro*. Violin plot showing HIF1α and active β-catenin MFI in WT or Tcf7l2^cKO^ AM, stimulated with Poly(I:C) for 24h *in vitro*. Scale bar 10 μm. B. Bar graph showing indicated gene expression of Wnt pathway in sorted GFP^+^ AM transduced with Adv-Control or Adv-Tcf7l2. C. Bar graph showing gene expression of indicated Wnt ligands in sorted GFP^+^ AMs transduced with Adv-Control or Adv-Tcf7l2. D. Bar graph showing gene expression of indicated Wnt ligands in sorted GFP^+^ AMs transduced with Adv-Control or Adv-Tcf7l2. E. Graph showing oxygen consumption rate in Tcf7l2^fl/fl^ or Tcf7l2^cKO^ AM treated with HIF-1α and active β-catenin for 24h *in vitro*. Bar graph showing basal and maximum oxygen consummation rate. F. Bar graph ChIP analysis of β-catenin and TCF4 binding to *Enol*, and *Hk2* in WT AMs treated with Wnt activator. G. Bar graph ChIP analysis of β-catenin and TCF4 binding to *Ndufs1*, and *Ndufa4* in WT AMs treated with Wnt activator. Data are representative of at least two independent experiments with similar results. *p*-values are represented as ns = non-significant, \**p*< 0.05, \*\**p*< 0.01, \*\*\*\**p*< 0.001.

**Fig. S4.**
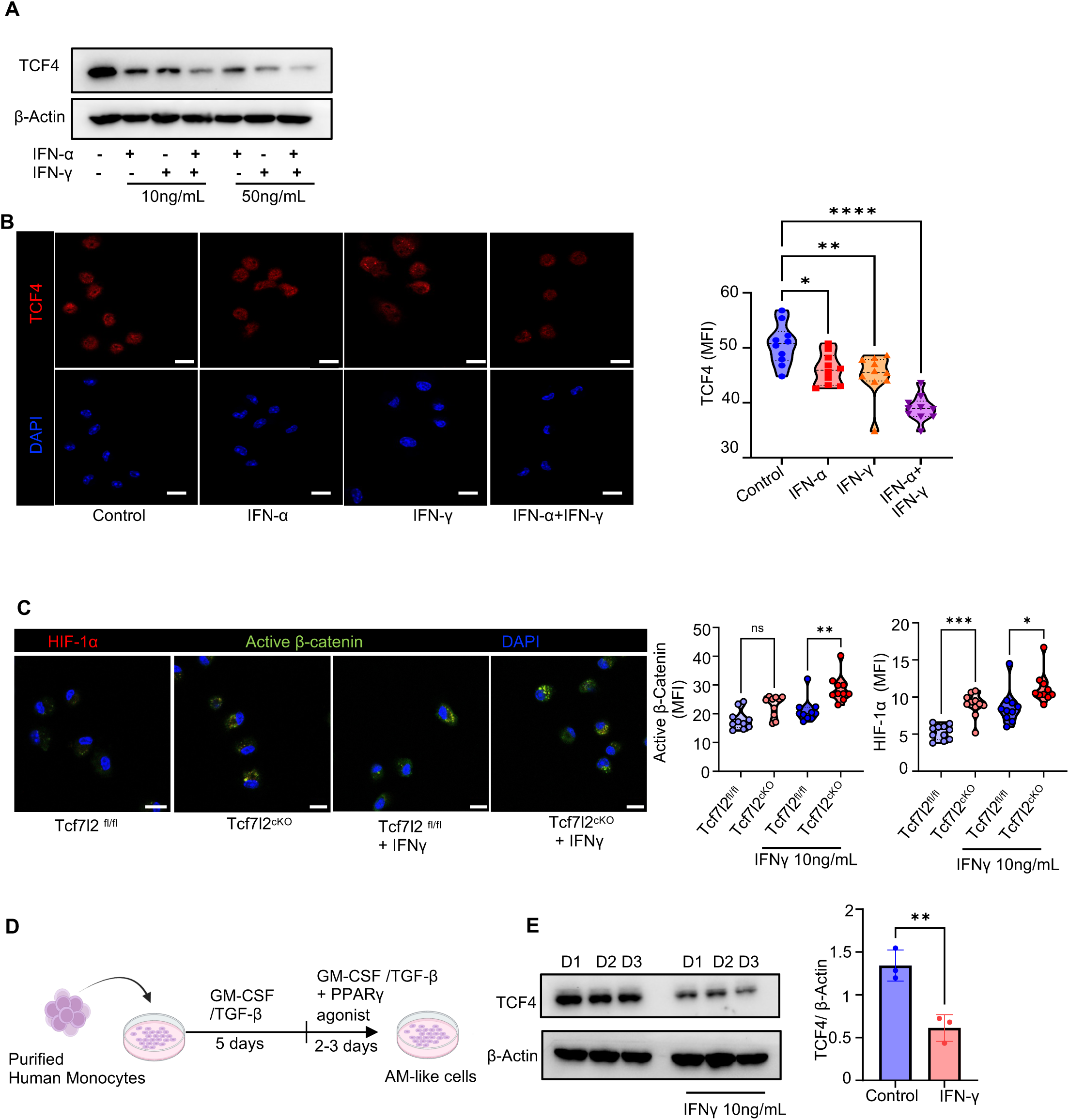
IFN signaling regulates TCF4 expression. A. Western Blot showing TCF4 expression in WT AMs stimulated with IFN-α and/or IFN-γ for 24h *in vitro*. B. Representative confocal images in WT AMs stimulated with IFN-α and/or IFN-γ. Right violin plot showing quantification of TCF4 under IFN stimulation. C. Representative confocal images in WT AM stimulated with IFN-α and/or IFN-γ. Right violin plot showing quantification of HIF-1α and active β-catenin under IFN stimulation D. Schematic diagram of the differentiation of purified monocytes from human donors to AM-like macrophages. E. Western blot showing TCF4 expression in monocytes derived AMs from 3 different individuals, donor (D)1-3, under stimulation of IFNγ for 24h. Right graph showing quantification of TCF4 expression under stimulation of IFNγ for 24h. Data are representative of at least two independent experiments with similar results, *p*-values are represented as ns = non-significant, \**p*< 0.05, \*\**p*< 0.01, \*\*\**p*< 0.005, \*\*\*\**p*< 0.001.

**Fig. S5.**
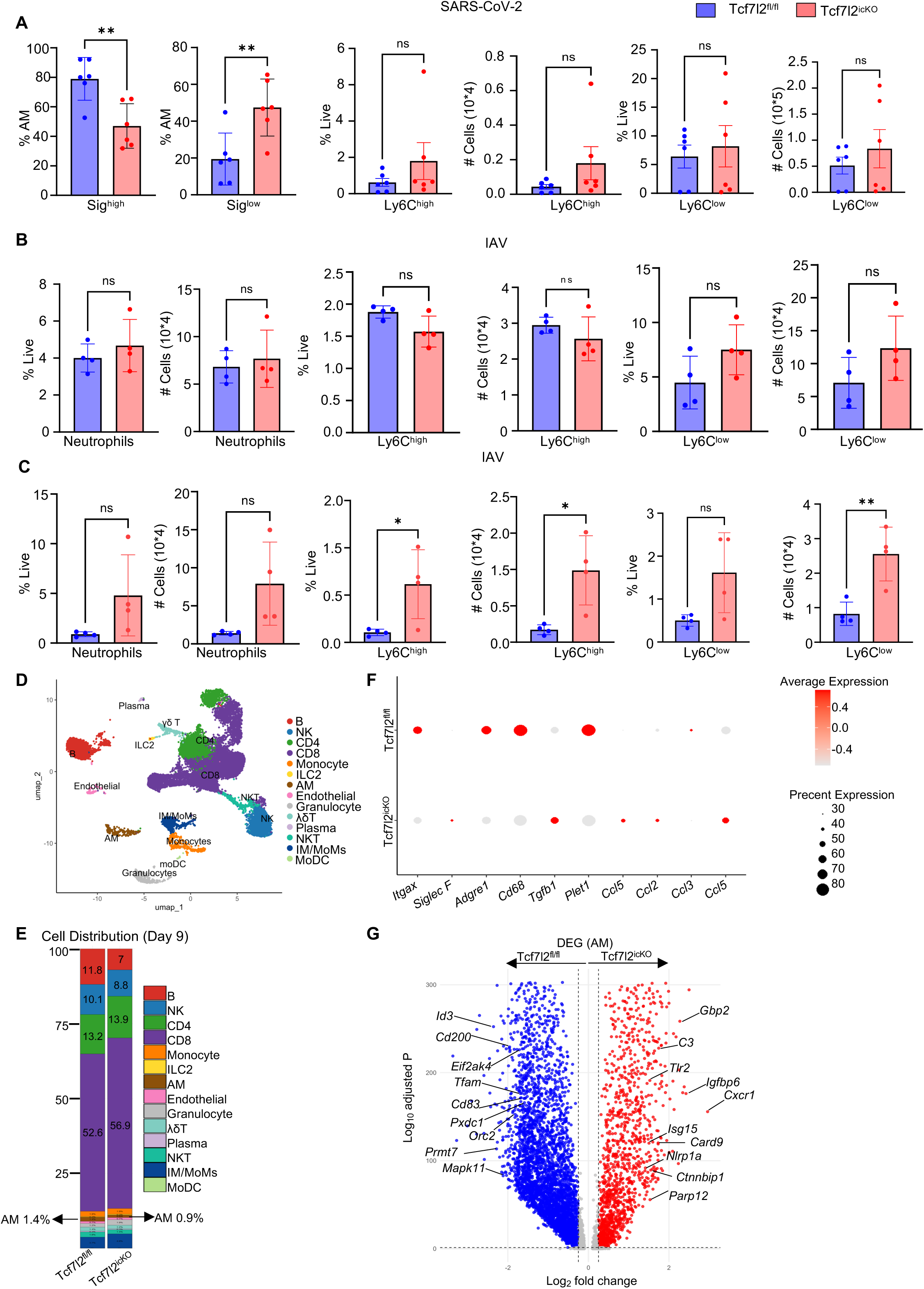
TCF4 deficiency results in poor AM recovery from SARS-COV-2 or IAV infection. A. Tcf7l2^fl/fl^ or Tcf7l2^icKO^ mice were injected with tamoxifen and then infected with SARS-CoV-2 MA30 as in Fig. 5A. Bar graph showing frequency and count of Siglec F^high^ AMs, Siglec F^low^ AMs, Ly6C^high^ and Ly6C^low^ monocytes in SARS-CoV-2 infected Tcf7l2^fl/fl^ (n=6) or Tcf7l2^icKO^ (n=6) mouse BAL at 10 d.p.i. B. Tcf7l2^fl/fl^ or Tcf7l2^icKO^ mice were injected with tamoxifen and then infected with IAV as in Fig. 5E. Bar graph showing frequency and count of neutrophil, Ly6C^high^ monocytes, and Ly6C^low^ monocyte, in IAV infected Tcf7l2^fl/fl^ (n=4) or Tcf7l2^icKO^ (n=4) mouse BAL at 14 d.p.i. C. Tcf7l2^fl/fl^ or Tcf7l2^icKO^ mice were infected with IAV and then injected with tamoxifen as in Fig. 5I. Bar graph showing frequency and count of neutrophil, Ly6C^high^ monocytes, and Ly6C^low^ monocyte in IAV infected Tcf7l2^fl/fl^ (n=4) and Tcf7l2^icKO^ (n=4) mouse BAL at 14 d.p.i. D. UMAP analysis showing different cell population in lung from IAV infected Tcf7l2^fl/fl^ or Tcf7l2^icKO^ mice at day 9 with scRNAseq analysis as in Fig. 5 I-P. E. Bar graph showing cell distribution in lung from IAV infected Tcf7l2^fl/fl^ or Tcf7l2^icKO^ mice at day 9. F. Dot plot showing enriched genes in AM from IAV infected Tcf7l2^fl/fl^ or Tcf7l2^icKO^ mice at day 9. G. Volcano plot showing DEG in AM from IAV infected Tcf7l2^fl/fl^ or Tcf7l2^icKO^ mice at day 9. Data are representative of at least two independent experiments with similar results except D-G, *p*-values are represented as ns = non-significant \**p*< 0.05, \*\**p*< 0.01.

**Fig. S6.**
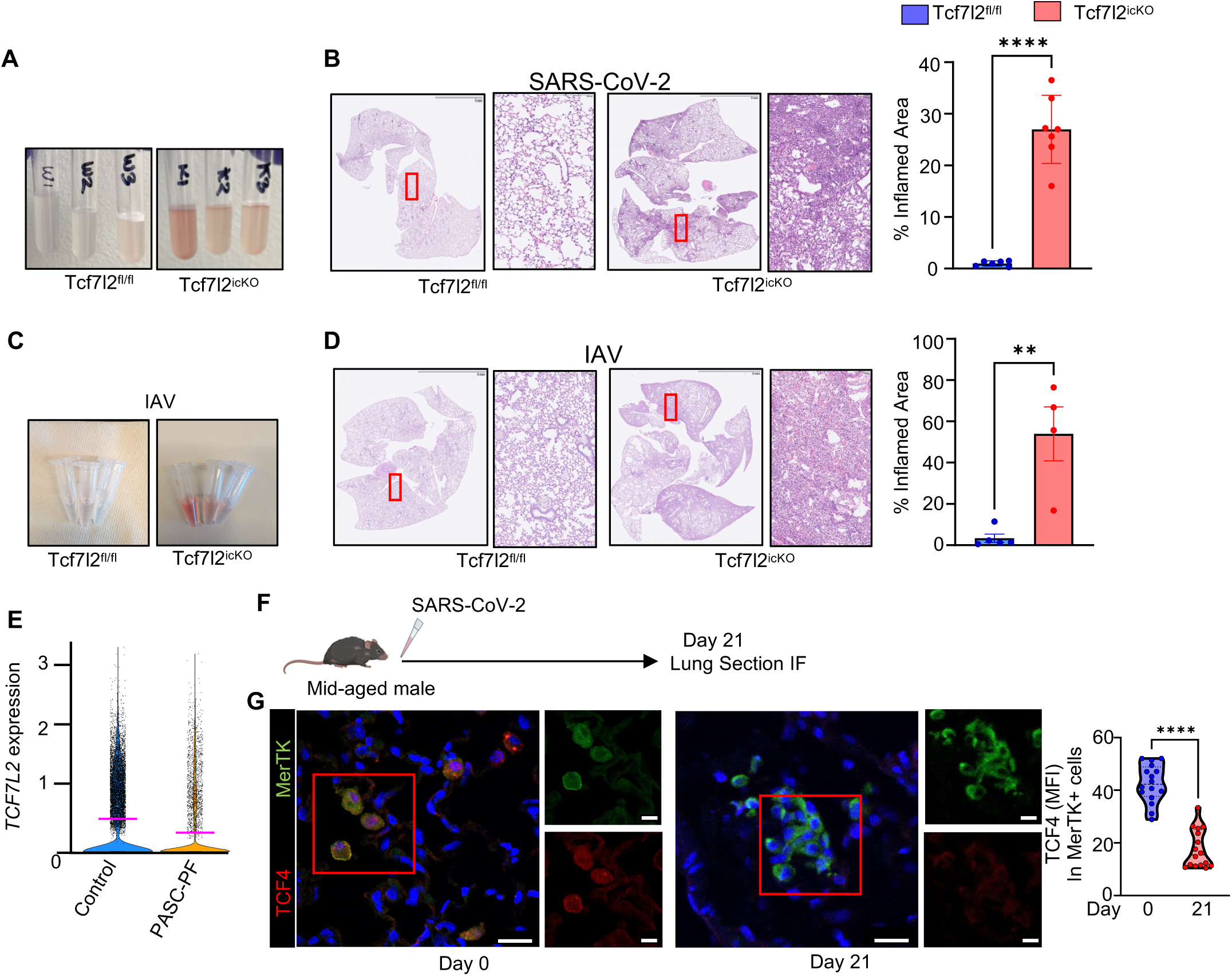
Myeloid TCF4 deletion results in chronic lung sequelae post SARS-CoV-2 or IAV infection. A. BAL image taken from Tcf7l2^fl/fl^ or Tcf7l2^icKO^ on day 21 after SARS-CoV-2 infection. B. Representative H&E of lungs Tcf7l2^fl/fl^ or Tcf7l2^icKO^ mice after SARS-CoV-2 infection. Bar graph showing % disrupted area in the lung of Tcf7l2^fl/fl^ or Tcf7l2^icKO^. C. BAL image taken from Tcf7l2^fl/fl^ or Tcf7l2^icKO^ on 42 days post IAV infection. D. Representative H&E of lungs Tcf7l2^fl/fl^ or Tcf7l2^icKO^ mice after IAV infection at 42 d.p.i. Bar graph showing % disrupted area in the lung of Tcf7l2^fl/fl^ or Tcf7l2^icKO^. E. Violin plot showing *TCF7L2* expression in AMs from healthy controls and PASC-PF patients from publicly available dataset (GSE224955). F. Schematic diagram showing the model of post-acute sequelae after SARS-CoV-2 virus infection in middle aged male WT mice. G. Representative confocal microscopy IF images of MerTK and TCF4 staining in lung samples from naïve and SARS-CoV-2 infected mid-aged male mice at 21 d.p.i. Violin plot showing quantification of TCF4 MFI in MerTK^+^ cells. Data are representative of at least two independent experiments with similar results, *p*-values are represented as \**p*< 0.05, \*\**p*< 0.01, \*\*\*\**p*< 0.001.

**Fig. S7.**
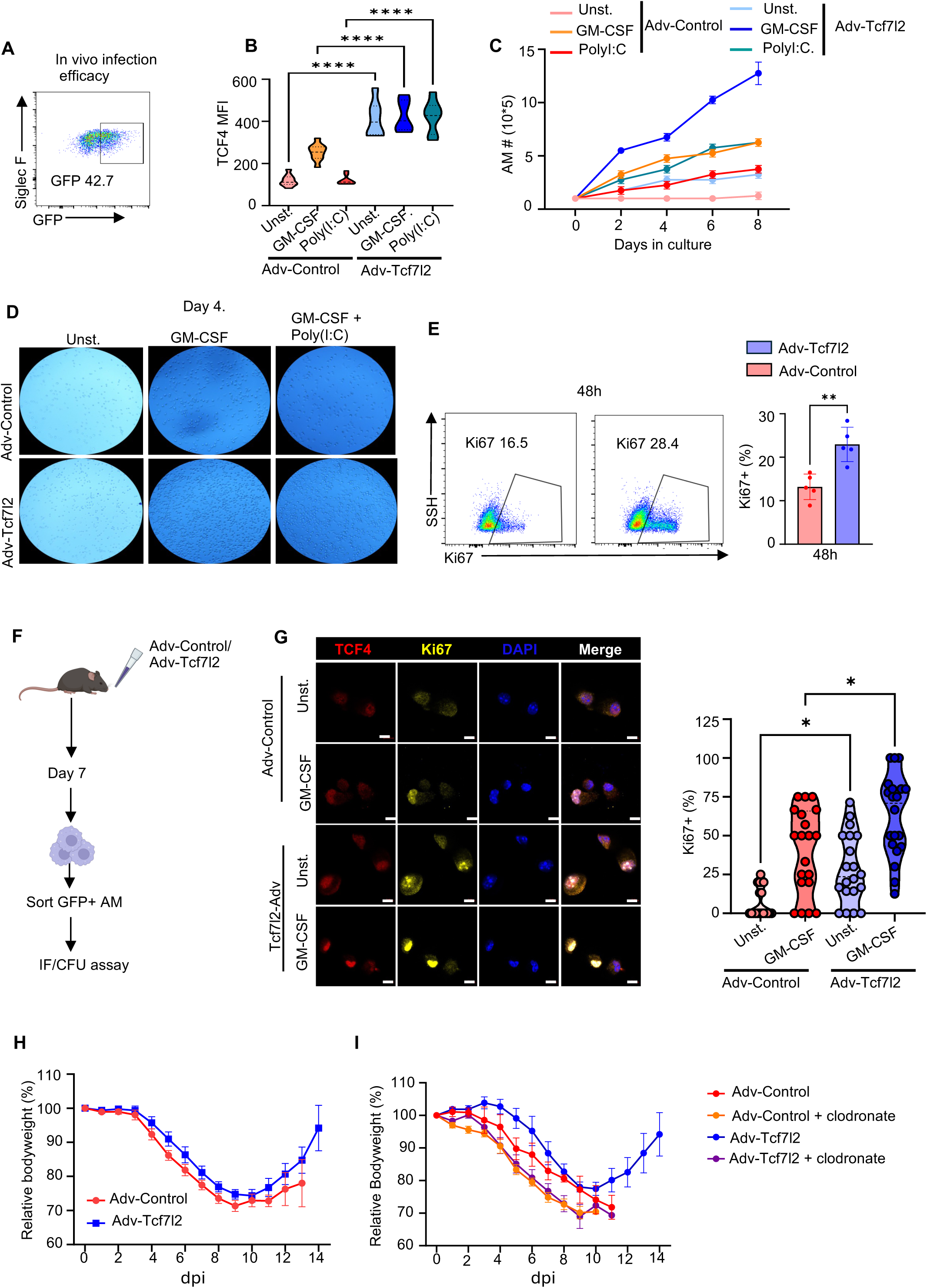
Enforced TCF4 expression promotes AM proliferation and protects from lethal infection. A. *In vivo* transduction efficiency of Adenovirus in WT mice as measured by GFP expression. B. Violin plot showing TCF4 expression in GFP+ AM transduced with Adv-Control and Adv-Tcf7l2 for 24h *in vitro*. C. Line graph showing total numbers of AMs transduced with Adv-Control or Adv-Tcf7l2 and culture with or without Poly(I:C) *in vitro*. D. Representative microscopy images for WT AMs on day 4. AM were transduced with Adv-Control or Adv-Tcf7l2 and culture with or without Poly(I:C) *in vitro*. E. Flow cytometry plot showing frequency of Ki67^+^ in WT AMs transduced with Adv-Control or Adv-Tcf7l2 *in vitro* for 48h. Right bar showing % Ki67^+^ in WT AMs transduced with Adv-Control and Adv-Tcf7l2 *in vitro* for 48h. F. Schematic for infection of WT mice with Adv-Control or Adv-Tcf7l2. On day 7 GFP^+^ AM were sorted and performed IF or CFU assay. GFP+ AM were also cultured *in vitro* with or without GM-CSF treatment. G. Representative confocal images of sorted GFP^+^ AMs culture with or without GM-CSF treatment for 48h and stained with TCF4 and Ki67. Scale bar 5 μm. Right bar showing quantification of Ki67^+^ AMs after Adv-Control or Adv-Tcf7l2 transduction for 48h *in vitro*. H. Relative percentages of weight loss of WT mice transduced with Adv-Control (n=13) or Adv-Tcf7l2 (n=14). The mice were later infected with lethal dose of IAV as in Fig. 7H. I. Relative percentages of weight loss of WT mice transduced with Adv-Control or Adv-Tcf7l2. AMs were depleted using clodronate in both groups after Adv-Control or Adv-Tcf7l2 transduction. The mice were later infected with lethal dose of IAV as in Fig. 7J. Data are representative of at least two independent experiments with similar results. *p*-values are represented as \**p*< 0.05, \*\**p*< 0.01, \*\*\*\**p*< 0.001.

**Fig. S8.**
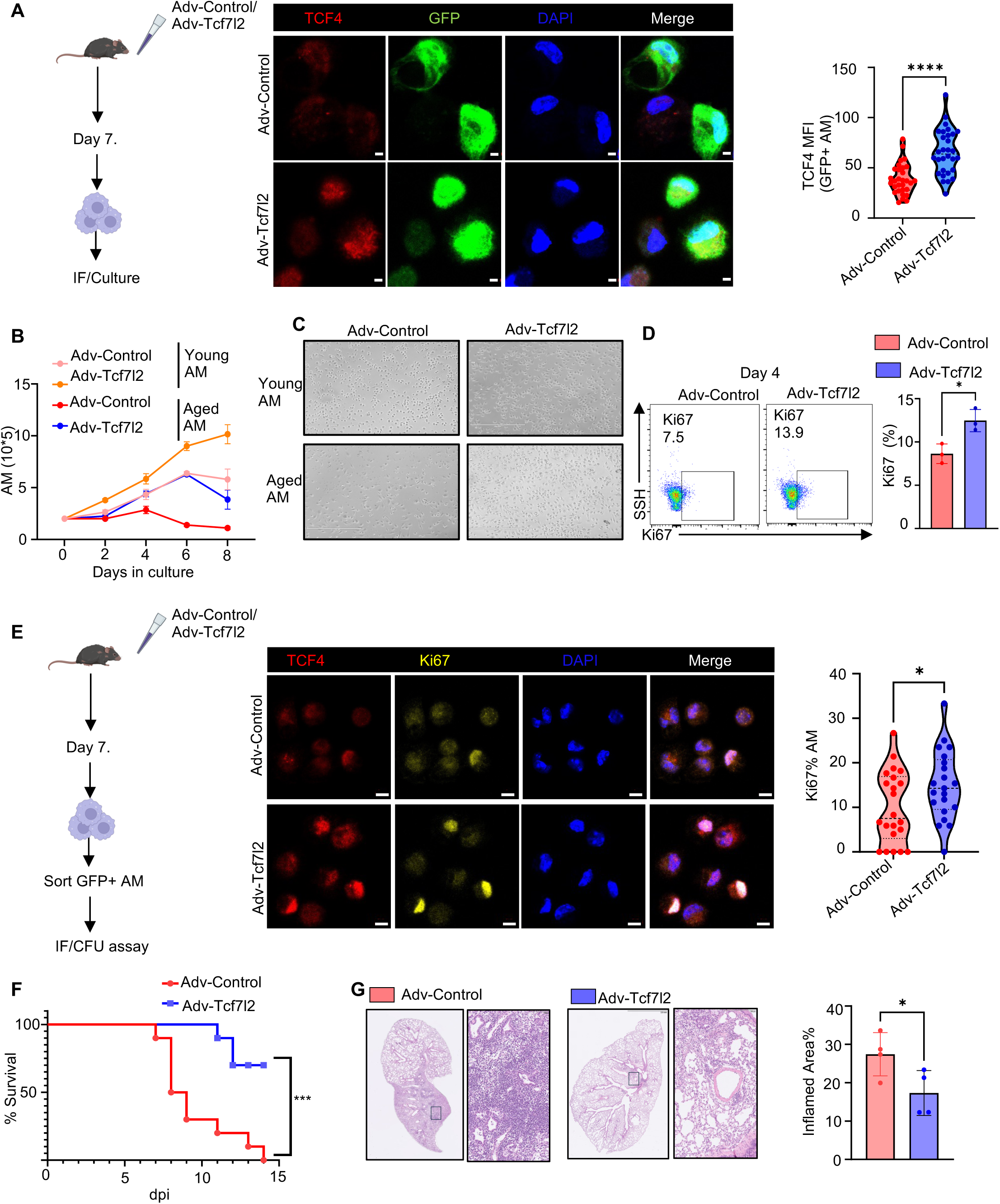
TCF4 over-expression dampens age-associated decline of AM proliferation and increases host resistance to severe infection during aging. A. Schematic of transduction of aged mice with Adv-Control or Adv-Tcf7l2. AM were isolated at day 7 after adenovirus inoculation for culture and IF imaging. Representative confocal images for TCF4 and GFP in WT aged AM transduced with Adv-Control or Adv-Tcf7l2. Violin plot showing TCF4 expression in GFP^+^ AMs. B. The line graph showing total number of *in vitro* cultured AMs derived from young or aged mice after transduction of Adv-Control or Adv-Tcf7l2. C. Representative microscopic images showing total number of AMs derived from young or aged mice after transduction with Adv-Control or Adv-Tcf7l2. D. Flow cytometry plot showing frequency of Ki67^+^ in aged AMs after transduction of Adv-Control or Adv-Tcf7l2 at day 4 post *in vitro* culture. Right bar graph showing frequency of Ki67^+^ in AMs. Bar graph showing % Ki67^+^ AM in a field of WT AMs after transduction of Adv-Control or Adv-Tcf7l2 day 4 *in-vitro*. E. Schematic for infection of aged mice with Adv-Control or Adv-Tcf7l2. At day 7 GFP+ AM were sorted and cultured *in vitro* with GM-CSF treatment. Right representative confocal images for TCF4 and Ki67 in GFP+ sorted AMs. Violin plot showing % of Ki67^+^ in sorted GFP+ AM. F. Survival plot in aged WT mice transduced Adv-Control (n=10) or Adv-Tcf7l2 (n-10) and then infected with IAV (150PFU). G. Representative image for H&E on lung samples isolated on day 60 from IAV-infected aged mice that were transduced with Adv-Control (n=5) or Adv-Tcf7l2 (n=5) as in Fig. 8H. Bar graph on right showing % disrupted area in the lung. Data are representative of at least two independent experiments with similar results. *p*-values are represented as \**p*< 0.05, \*\*\**p*< 0.005, \*\*\*\**p*< 0.001.

